# Mycobacterial DnaQ is an Alternative Proofreader Ensuring DNA Replication Fidelity

**DOI:** 10.1101/2023.10.24.563508

**Authors:** Ming-Zhi Deng, Qingyun Liu, Shu-Jun Cui, Han Fu, Mingyu Gan, Yuan-Yuan Xu, Xia Cai, Wei Sha, Guo-Ping Zhao, Sarah M. Fortune, Liang-Dong Lyu

## Abstract

Remove of mis-incorporated nucleotides ensures replicative fidelity. Although the ε-exonuclease DnaQ is a well-established proofreader in the model organism *Escherichia coli*, proofreading in mycobacteria relies on the polymerase and histidinol phosphatase (PHP) domain of replicative polymerase despite the presence of an alternative DnaQ homolog. Here, we show that depletion of DnaQ in *Mycolicibacterium smegmatis* results in increased mutation rate, leading to AT-biased mutagenesis and elevated insertions/deletions in homopolymer tract. We demonstrated that mycobacterial DnaQ binds to the β-clamp and functions synergistically with the PHP domain to correct replication errors. Further, we found that the mycobacterial DnaQ sustains replicative fidelity upon chromosome topological stress. Intriguingly, we showed that a naturally evolved DnaQ variant prevalent in clinical *Mycobacterium tuberculosis* isolates enables hypermutability and is associated with extensive drug resistance. These results collectively establish that the alternative DnaQ functions in proofreading, and thus reveal that mycobacteria deploy two proofreaders to maintain replicative fidelity.

## Introduction

The emergence of drug-resistant *Mycobacterium tuberculosis* (*Mtb*) poses a great challenge to the global control of tuberculosis, which kills 1.5 million people annually^1^. Unlike many other bacteria, in which the development of drug resistance is typically driven by horizontal gene transfer (HGT), genome rearrangement and mutation, all drug resistances characterized in *Mtb* are the result of chromosomal mutations, indicating that the mutational capacity of *Mtb* is one of the key mechanisms underpinning its *de novo* generation of drug resistance^2,3^.

Nucleotide mispairing occurred during DNA replication is a major source of mutagenesis. In the model organism *Escherichia coli*, the fidelity of DNA replication is ensured by the base selection via DNA polymerase *α* subunit (DnaE), the proofreading by the ε subunit 3’-5’ exonuclease (DnaQ) and the post-replicative mismatch repair (MMR) committed by a separate enzyme system (MutHLS)^3–5^. Proofreading functions as a double-checking point that removes the mis-incorporated nucleotide *via* 3’-5’ exonuclease activity^6^. In *E. coli*, the naturally occurred mutants defective in proofreading activity can give rise to 100- to 1000-fold increase of mutation rate and can accelerate the evolving of drug resistance^7,8^. In addition, deletion of *E. coli dnaQ* also results in growth defect, insufficient polymerization capacity and constitutive SOS phenotype^6,9,10^.

The *E. coli* DnaQ is a single-domain protein containing a DEDDh-family exonuclease domain followed immediately by a clamp-binding motif (CBM)^11^. Mycobacteria encode two DnaQ homologs which share considerable similarity to the *E. coli* DnaQ in their N-terminal exonuclease domain but generally differ in size and domain organization as a whole (**Supplementary Fig. 1**). The annotated DnaQ contains an extra C-terminal domain of 86 residues homologous to the human breast cancer suppressor protein C-terminal domain (BRCT), which functions as a protein-protein interaction module found in large variety of proteins involved in DNA replication, repair, and recombination^12^. The second potential DnaQ homolog (annotated as a hypothetical protein) has a larger C-terminal region of ~400 residues which exhibits high similarity to the endonuclease domain of the nucleotide excision repair protein UvrC. However, despite the *de facto* exonuclease activity of *Mtb* DnaQ *in vitro*, previous fluctuation analysis results showed that deletion of the annotated *dnaQ* (*Rv3711c*) in *Mtb* or deletion of both two *dnaQ* homolog genes (*Ms6275* and *Ms4259*) in *Mycolicibacterium smegmatis* (*Msm*) did not lead to the “expected” mutator phenotype^11^. This functional departure from that of the *E. coli* DnaQ led to the identification of a novel proofreading activity mediated by the polymerase and histidinol phosphatase (PHP) domain of the *dnaE1*-encoded replicative polymerase. Together, these findings demonstrate that the mycobacterial DnaQ is an alternative exonuclease that is structurally and functionally distinct from the *E. coli* DnaQ.

Phylogenetic and structural analyses suggested that the PHP domain is the most common replicative exonuclease in the bacterial kingdom^11,13,14^. Intriguingly, although the presence of active PHP domain appears to be mutually exclusive with the presence of an *E. coli*-like ε-exonuclease (DnaQ), it extensively co-exists with the alternative DnaQ^11^. However, the biological function of this kind of alternative exonuclease remains unclear^15^. Here, we show that the mycobacterial DnaQ functions in proofreading and exhibits additional role in maintenance of genomic GC content and correcting of replication errors upon chromosome topological stress. Using the mutation accumulation (MA) assay combined with whole genome sequencing, we demonstrated that deletion of *dnaQ* results in 2.4-fold increase of spontaneous mutation rate in *Msm*, leading to a mutational bias for AT and increased insertions/deletions in homopolymer tract. We demonstrate that mycobacterial DnaQ functions synergistically with PHP domain-mediated proofreading and physiologically associates with the β-clamp. Moreover, loss of *dnaQ* results in replication fork dysfunction upon topological stress, leading to attenuated growth and increased mutagenesis. Importantly, through analyzing the sequence polymorphism from 51,229 *Mtb* clinical isolates, we provide real-world evidence that *dnaQ* was subjected to positive selection in some sublineages and had functional effect on mutagenesis. Particularly, we show that a a naturally evolved DnaQ variant prevalent in lineage 4.3 (16.5%) enables *Mtb* hypermutability and is associated with extensive drug resistance, suggesting that the mutator-like evolution trajectory may exist during *Mtb*’s adaptation. These findings collectively establish that mycobacterial DnaQ is an alternative proofreader and provide a new action model of DNA replication proofreading deploying two proofreaders (PHP and DnaQ), which may have broad implication with respect to DNA metabolism.

## Results

### Deletion of *dnaQ* results in increased mutation rate in *Msm*

To investigate whether mycobacterial *dnaQ* contributes to the maintenance of DNA replication fidelity, we constructed the null mutant strain *ΔdnaQ, ΔMs4259* (which encodes the second potential *dnaQ* homolog) and *ΔdnaQΔMs4259* in *Msm* and estimated the mutation rate by fluctuation analysis (**Fig.S2A**). Deletion of these genes did not affect bacterial growth in vitro (**Fig.S2B**). Consistent with previous study^11^, our results showed that these mutant strains did not exhibit significant increase of spontaneous mutation rate (defined by non-overlapping 95% confidence intervals [CI]) (**Fig. 1A** and **Table S1**). However, we found that deficiency in *dnaQ* increased the mutation frequency (number of mutants per cell plated in a single culture) of rifampicin resistance (Rif^R^) by 3-fold (*P*<0.05) and this phenotype can be complemented by expression of wild-type *dnaQ* from either *Msm* or *Mtb* strain H37Rv, but not by the expression of exonuclease activity deficient variant (Exo^−^) DnaQ[D28A/E30A/D112A] (**Fig. 1A**).

**Figure 1.**
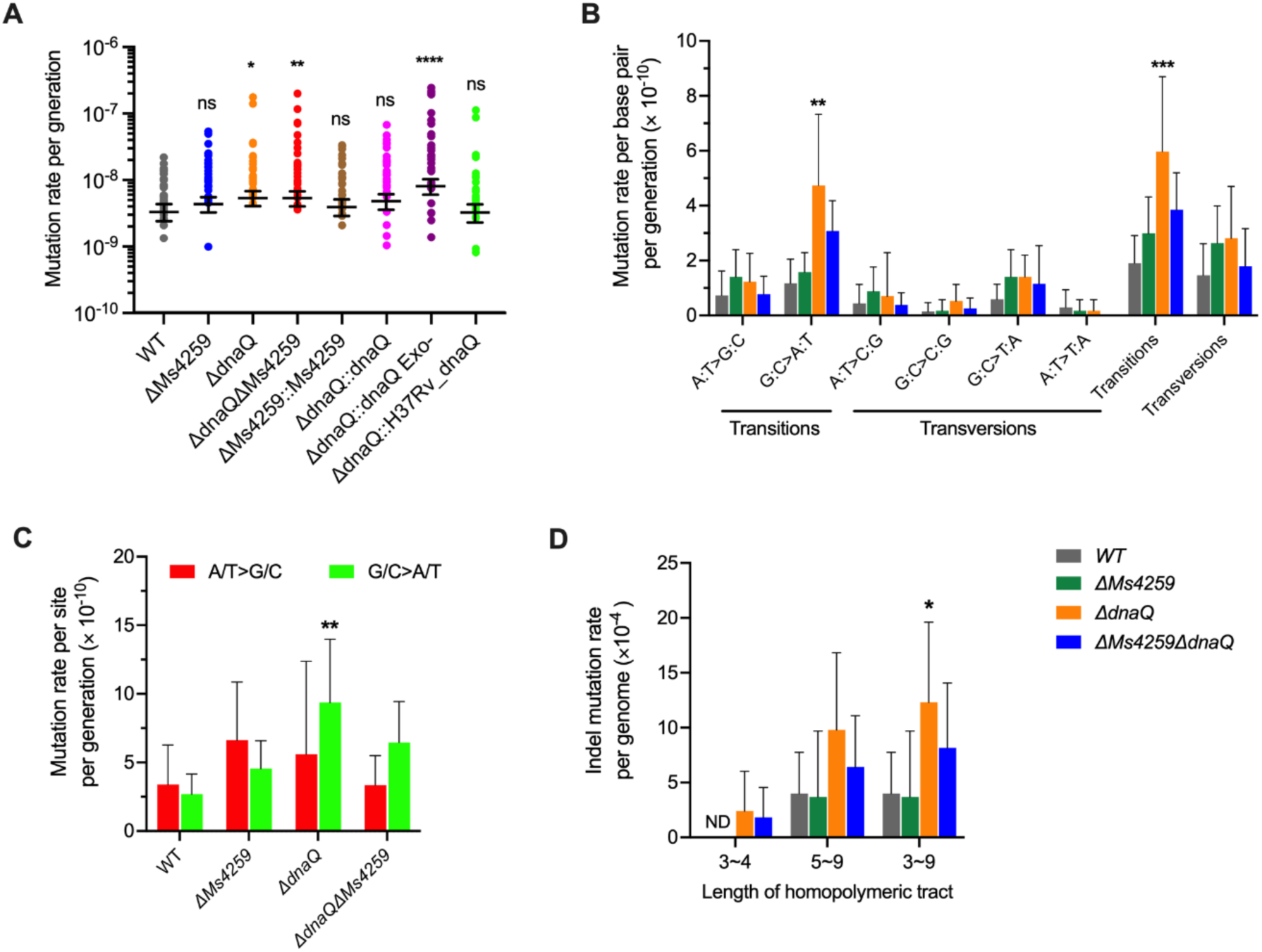
Deletion of *dnaQ* in *Msm* results in a mutational bias for AT and increased insertions/deletions in homopolymeric tract. (A) Fluctuation analysis in *Msm* strains. *dnaQ* Exo^−^, DnaQ[D28A/E30A/D112A] variant deficient in exonuclease activity; H37Rv_*dnaQ*, *dnaQ* allele from *Mtb* strain H37Rv. Circles represent mutant frequency (number of rifampicin-resistant mutants per cell plated). Estimated mutation rates (mutations conferring rifampicin resistance per generation) were shown as mean ± 95% CI. Significance was determined using a one-way analysis of variance test on log transformed values; \**P* < 0.05, \*\**P* < 0. 01, \*\*\*\**P* < 0.0001. (B-D) Mutational spectra identified from the MA assay. (B) Mutation rates of each of the BPS mutational spectra. (C) Mutation rates of the AT/>GC and G/C>AT events normalized to the genomic AT and GC content, respectively. (D) Mutation rates of the indel events occurred in homopolymeric tract. Data shown are mean ± 95% CI. \**P*<0.05, \*\**P*<0.01, \*\*\**P*<0.001 by the Mann–Whitney U test compared to the counterpart of WT.

Because the fluctuation analysis assay relies on the selection of mutant (*e.g*., Rif^R^) harboring mutations in a specific locus (*e.g*., *rpoB*), the estimated mutation rate may not be representative of the genome. In addition, because selection is likely to affect putatively neutral sites, this method may have significant bias on detection of mutational events^16,17^. To accurately determine the mutation rate, we next deployed the MA experiment combined with whole-genome sequencing. For each strain, 10~12 MA lines were initiated from a single colony (**Table 1**). Every 2 d, a single colony from each line was re-streaked onto a fresh 7H10 plate. In this process, the strong bottlenecks minimize the selective pressure, enabling mutations to accumulate in a neutral unbiased manner. The application of whole-genome sequencing to MA lines further ensures the detection of complete and nearly unbiased profile of mutational events^5,16,17^. The MA experiment was carried out for 100 days, which accounted for > 8150 generations for each strain (**Table 1**). Totally 44 MA lineages were successfully sequenced, with an average of 99.96% of the reference genome were covered with >10x reads (**Supplemental dataset 1**). Through identify the base pair substitutions (BPSs) and insertions/deletions (indels, defined as insertions or deletions of ≤ 30 nucleotides) (**Supplemental dataset 1**), we found that the basal mutation rate for *Msm* wild type (WT) was 4.88 × 10^−10^ mutations per base pair per generation (95% CI: [2.97-6.78] × 10^−10^), consistent with the estimates of previously studies (95% CI: [4.52-5.27] × 10^−10^)^11,18^. Among the mutant strains, the mutation rate of the *ΔdnaQ* strain was 11.68 × 10^−10^ mutations per base pair per generation (95% CI: [7.05-16.31] × 10^−10^), a 2.4-fold increase over the WT (**Table 1**). Deletion of the *dnaQ* homolog encoded by *Ms4259* either in the *dnaQ*^+^ (*ΔMs4259*) or in the *ΔdnaQ* strain (*ΔdnaQΔMs4259*) resulted in moderate increase (~1.4-fold) of mutation rate (**Table 1**).

**Table 1.** Deletion of *dnaQ* results in a mutator phenotype in mutation accumulation assay.

| Strain | No.<br>of BPSs | No.<br>of indels | No.<br>of lines | Generations<br>per line | Total No. of<br>generations | Mutation rate<br>per base pair ± 95% CI<br>($\times 10^{-10}$) | Mutation rate<br>per genome ± 95% CI<br>($\times 10^{-3}$) |
| --- | --- | --- | --- | --- | --- | --- | --- |
| *WT* | 23 | 10 | 12 | 815 | 9780 | $4.88 \pm 1.90$ | $3.39 \pm 1.29$ |
| $\Delta dnaQ$ | 50 | 16 | 10 | 815 | 8150 | $11.68 \pm 4.63$ | $8.08 \pm 3.14$ |
| $\Delta Ms4259$ | 32 | 7 | 10 | 815 | 8150 | $6.83 \pm 1.95$ | $4.78 \pm 1.40$ |
| $\Delta Ms4259 \Delta dnaQ$ | 44 | 11 | 12 | 930 | 11160 | $7.09 \pm 2.36$ | $4.94 \pm 1.62$ |

Mutation rates were estimated by mutant accumulation assay. Mutations are classified according to base-pair substitutions (BPSs) or indels (defined here as insertions or deletions of $\leq 30$ nucleotides). CI, confidence intervals.

### Depletion of DnaQ leads to a mutation bias for AT and increased indels in homopolymeric tract

In WT, we identified totally 23 BPSs across the 12 lineages, showing a transition/transversion ratio of 1.30 (13/10) (**Table S2**), which is very similar to the previous observation (1.48)^18^. However, this ratio was elevated to 2.13 (34/16) in the *ΔdnaQ* mutant, yielding a 3.1-fold (*P*<0.001) increase of transition mutation rate over the WT (**Fig. 1B** and **Table S3**). Among the transitions, the G:C>A:T mutation rate significantly increased 4.2-fold (*P*<0.01) in the *ΔdnaQ* mutant compared to that of the WT (**Fig. 1B** and **Table S3**). Given that the G:C>A:T mutation has a great influence on genomic GC content during long-term evolution ^19^, we analyzed the direction of mutation bias (**Fig. 1C** and **Table S4**). In WT, the mutation rate of A/T>G/C (*i.e.* A to G or C, T to G or C) and G/C>A/T normalized to the genomic GC or AT content are 3.40 × 10^−10^ and 2.69 × 10^−10^, respectively, thus exhibiting a mutation bias for GC (expected GC content is 55.8%, the actual genomic GC content of *Msm* is 65.6%), which is consistent to previous MA study (expected GC content was 58.2%)^18^. In contrast, while the mutation rate of A/T>G/C remained unchanged in the *ΔdnaQ* mutant, the G/C>A/T mutation rate (95% CI: [4.75-13.98] × 10^−10^) significantly increased 3.5-fold (*P*<0.01), resulting in a striking bias for AT mutation (expected GC content of 37.4%) (**Fig. 1C**). Deletion of the *dnaQ* homolog (*Ms4259*) in the *ΔdnaQ* background (*ΔdnaQΔMs4259*) resulted in similar mutation spectra and mutational bias for AT as that of the *ΔdnaQ* mutant, however, these effects were not observed in the *ΔMs4259* mutant (**Fig.1B-C**, **Table S3 and Table S4**). Together, these results indicate that mycobacterial *dnaQ* implicates in maintenance of genomic GC content. Of note, the mutation spectrum of the mycobacterial *ΔdnaQ* mutant is quite distinct from that of the *E. coli* mutant deficient in *dnaQ*, which showed a mutation bias for transversion and no effect on GC content^20^.

Our MA analysis identified totally 44 indels 1-30 bp in length (**Table 1** and **Supplemental dataset 1**), and all strains exhibited a mutational bias for insertion events (**Table S2**). The indel rate of WT (1.48 × 10^−10^) is very similar to the previous estimate (1.27 × 10^−10^)^18^. While the *ΔdnaQ* mutant showed a 1.9-fold increase of indel mutation rate (**Table S5**), the rate in the *ΔMs4259* strain exhibited a slightly reduction, suggesting a functional departure between DnaQ and Ms4259. Strikingly, the indel events in the *ΔdnaQ*, *ΔMs4259* and *ΔdnaQΔMs4259* mutants occurred more frequently in the lagging strands (62.5%, 71% and 73%, respectively) than that of the WT (40%) (**Supplemental dataset 1**). In the WT and the *ΔMs4259* strain, about 40% of the indels occurred in homopolymer tract. However, these fractions were increased to 63% and 82% in the *ΔdnaQ* mutant and the *ΔdnaQΔMs4259* mutant, respectively (**Table S5**). Accordingly, the mutation rate of indels occurred in homopolymeric tract (almost G/C tract) was significantly increased 3-fold (*P*<0.05) in the *ΔdnaQ* mutant compared to the WT (**Fig. 1D** and **Table S5**). Moreover, indels events occurred in homopolymeric tract less than 5 nucleotides in length were exclusively observed in the *ΔdnaQ* mutant and the *ΔdnaQΔMs4259* mutant (**Fig. 1D** and **Table S6**). Given that indel events in the homopolymeric tract is mainly caused by strand slippage of replicative DNA polymerase^21^, these results indicate a role of mycobacterial *dnaQ* in DNA replication. Of note, recent studies showed that the indel events occurred in homopolymeric tract may have a strong impact on *Mtb*’s pathogenesis and drug tolerance^22–24^.

Collectively, the MA results established that mycobacterial *dnaQ* contributes to the maintenance of genetic information. Besides, our data indicated that the second potential DnaQ homolog encoded by *Ms4259* exhibits distinctive mutational phenotypes from the annotated *Msm* DnaQ. Our results also showed that the *ΔMs4259* mutant exhibited a mutation bias for noncoding region (account for 31% of the BPSs) (**Table S2**), which is significantly different from that of the *ΔdnaQ* mutant (8%) and from the expected value of 10% (*χ^2^*=16.6, *P*<0.01). Therefore, our data indicate that the second potential DnaQ homolog encoded in mycobacteria is functionally distinct from the annotated DnaQ.

### Mycobacterial DnaQ specifically corrects DNA replication errors

Spontaneous mutations can arise through replication errors or as a consequence of intrinsic DNA damage^17^. To determine whether the anti-mutational role of mycobacterial *dnaQ* is attributable to the correction of DNA replication errors, we performed fluctuation analysis and compared the mutational rate of WT and the *ΔdnaQ* mutant expressing DanE1[D228N] variant which is deficient in PHP domain-mediated proofreading^11^. Expression of DanE1[D228N] did not affect bacterial growth (**Fig. S2C**). While expression of wild-type *dnaE1* had no effect on the mutation rate in either the *dnaQ^+^* prototype strain (95% CI: [2.21-4.07] × 10^−9^) or the *ΔdnaQ* mutant strain (95% CI: [2.11-4.01] × 10^−9^), expression of DanE1[D228N] resulted in 9-fold increase of mutation rate in WT (95% CI: [2.21-3.41] × 10^−8^) and 17-fold in the *ΔdnaQ* mutant (95% CI: [4.18-5.95] × 10^−8^) (**Fig. 2A**). Therefore, the combination of *dnaQ* depletion and the deficiency in PHP proofreading activity results in a synergistical effect on mutagenesis, demonstrating that mycobacterial DnaQ participates in correction of replication errors.

**Figure 2.**
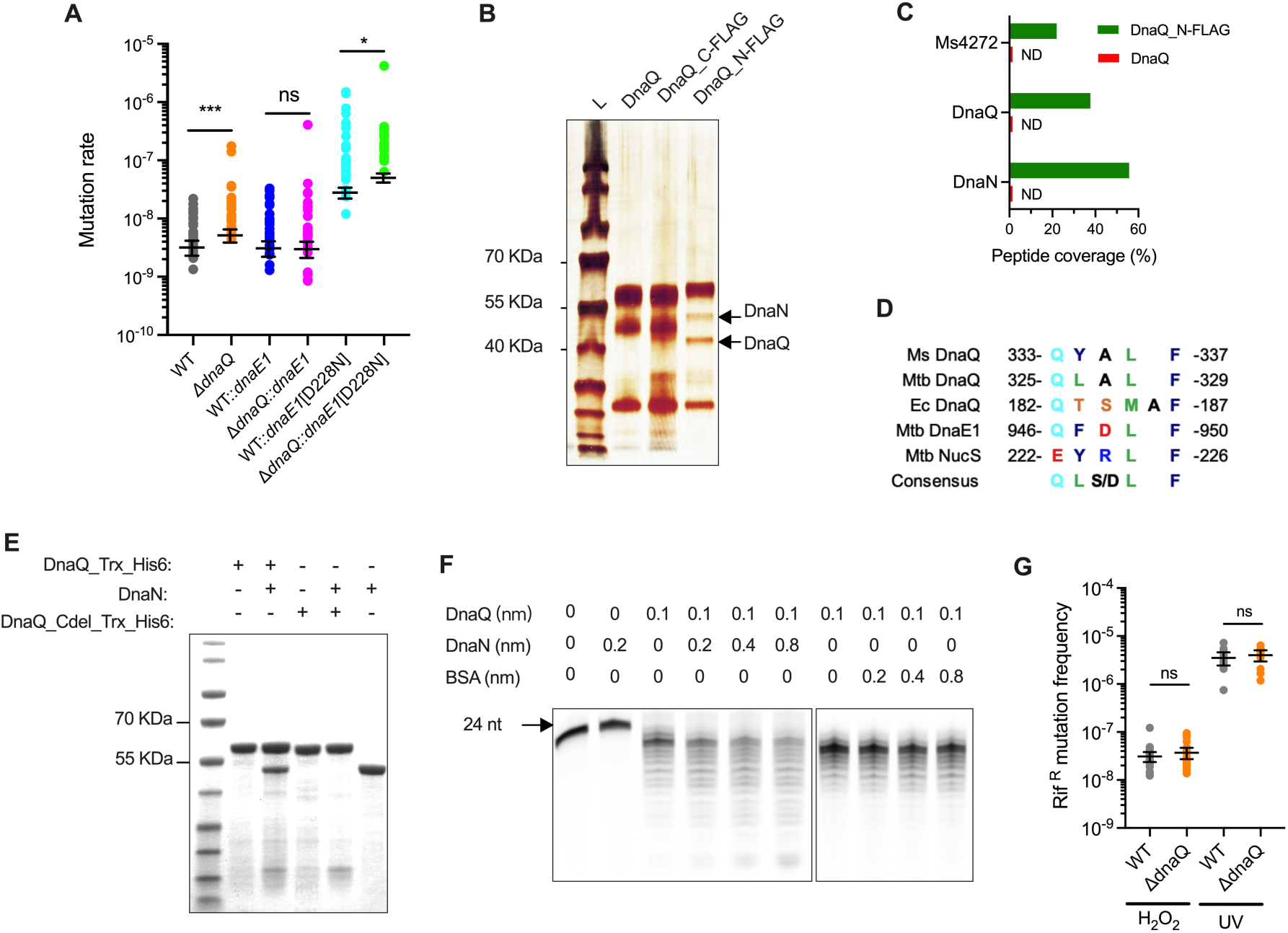
Mycobacterial DnaQ specifically corrects errors produced during DNA replication. (A) Fluctuation analysis in *Msm* strains expressing of wild-type *dnaE1* or allele encoding a mutant deficient in PHP-mediated proofreading activity (*dnaE1*[D228N]). Circles represent mutant frequency. Estimated mutation rates (mutations conferring rifampicin resistance per generation) were shown as mean ± 95% CI. Significance was determined by comparing strain pairs using an unpaired t test on log transformed values; \**P* < 0.05, \*\*\**P* < 0.001. (B) Immunoprecipitation eluates analyzed by silver staining. Immunoprecipitation experiments were performed with FLAG antibody in the *ΔdnaQ* mutant strain expressing either wild-type *dnaQ* (DnaQ) or alleles encoding DnaQ containing a FLAG tag at C terminal (DnaQ_C-FLAG) or N terminal (DnaQ_N-FLAG). Precipitated DnaQ and DnaN were identified by LC–MS/MS. (C) Peptide coverage of the precipitated proteins identified by LC–MS/MS. (D) Sequences of the clamp-binding motif (CBM) identified in mycobacterial DnaQ, DnaE1, NucS, and *E. coli* (Ec) DnaQ. Numbers indicate the peptide position. (E) Pull-down eluates analyzed by Coomassie blue staining. DnaQ_Trx_His6, DnaQ fused with a N-termial Trx domain containing 6×His tag; DnaQ_Cdel_Trx_His6, DnaQ_Trx_His6 with deletion of C-terminal CBM. DnaN, native DnaN without His tag. (F) DnaQ 3’-5’ exonuclease activity on single-strand DNA (ssDNA). Reactions were performed with a 24nt FAM-labeled ssDNA for 5 minutes. (G) Rif^R^ mutation frequencies of the indicated strains exposed to 5mM H_2_O_2_ or 25mJ/cm^2^ UV. Data shown are mean ± 95% CI.

To investigate whether DnaQ interacts with the replisome, we next performed in vivo immunoprecipitation using the *ΔdnaQ* mutant expressing DnaQ fusion protein containing a FLAG tag at either N-terminal (DnaQ_N-FLAG) or C-terminal (DnaQ_C-FLAG). Mass spectroscopy results demonstrated that immunoprecipitation of whole-cell lysates with FLAG antibodies coprecipitated DnaN and a protein with unknown function (*Ms4272*) from the strain expressing DnaQ_N-FLAG, but not from the lysate containing DnaQ_C-FLAG or untagged DnaQ (**Fig. 2B-C and Fig. S3A-C**). DnaN is the β subunit of DNA polymerase III holoenzyme, associates in pairs to form the β-clamp that encircles and slides along the DNA strands as replication proceeds. We found that mycobacterial DnaQ contains a highly conserved CBM Q[Y/L]ALF at its C-terminal proximity and the pulldown results demonstrated that this CBM is essential for the direct interaction between DnaQ and DnaN *in vitro*^25^ (**Fig. 2D-E**). Therefore, the inability of DnaQ_C-FLAG to coprecipitate DnaN may likely be due to the inaccessibility of CBM to DnaN caused by the fused tag. Further, our in vitro 3’-5’ exonuclease enzyme assay demonstrated that the presence of DnaN could increase the 3’-5’ exonuclease activity of DnaQ (**Fig. 2F and Fig. S3D**), suggesting that DnaQ biochemically prefers to act at the replication fork. Unlike *E. coli*, whereby the DnaQ interacts with both the β-clamp and the *α* subunit of DNA polymerase III (**Fig. S1B**)^26^, our results indicated that mycobacterial DnaQ did not form a stable complex with DnaE1, which are consistent with previous finding^11^.

In bacteria, the replication sliding clamp also recruits DNA repair proteins to the replication forks^25^.To assess whether mycobacterial DnaQ plays a role in DNA damage repair, we measured the stress-induced mutation frequency of rifampicin resistance (Rif^R^) in strains exposed to hydrogen peroxide (H_2_O_2_) or UV radiation, which are the most common source of endogenous and exogenous DNA damage, respectively^27^. Exposure of the WT to H_2_O_2_ and UV led to 4- and 90-fold increases of the Rif^R^ frequency, respectively (**Fig. 1A and 2G**). However, these stress-induced mutations were independent of *dnaQ*, as the *ΔdnaQ* mutant exhibited similar Rif^R^ frequency to WT (**Fig. 2G**). These results collectively indicate that mycobacterial DnaQ does not function in DNA damage repair.

### Depletion of DnaQ results in replication fork perturbation upon topological stress

The above results indicate that mycobacterial replisome may deploy two proofreaders (the PHP domain of DnaE1 and DnaQ) to correct replication errors. To further explore the physiological role of DnaQ, the additional proofreader, we performed transcriptional profiling and analyzed the DNA damage response (DDR) signature during normal growth. Mycobacterial cell deploys sophisticated regulatory systems that governs DDR against different types of DNA insults^28^. According to the previously published criteria (log_2_ fold change of ≥1.5, FDR<0.001)^28^, 46 DDR genes showed differential regulation (mostly upregulated *vs* the *dnaQ^+^* prototype strain) in the *ΔdnaQ* mutant, accounting for 40% of the differentially regulated genes (46/114) (**Fig. 3A, Supplementary dataset 2**). In contrast, only 3 DDR genes were upregulated in the *ΔMs4259* mutant strain. Strikingly, a dominant proportion (33/46) of the differentially expressed DDR genes in the *ΔdnaQ* mutant belongs to the PafBC regulon^28^, resulting in an enrichment over 29-fold (*P*<10^−15^, Fisher’s exact test) (**Fig. 3B and Fig. S4A**). These transcriptional signatures were further validated by qRT-PCR (**Fig. S4B**). Because the PafBC regulator responses specifically to quinolone antibiotics and replication fork perturbation^28^, as signified by the upregulation of the well-characterized recombinational fork-repair genes including *recBC*, *adnAB* and *sbcD* (**Fig. 3C**)^29–31^, these results demonstrate that DnaQ deficiency leads to dysfunction of replication fork during normal growth. In contrast to *E. coli dnaQ* mutant strain, which exhibited constitutive SOS phenotype and growth defect^9^, we did not observe apparent SOS signature in the *Msm ΔdnaQ* mutant during normal growth (**Supplementary dataset 2**). Further, our qRT-PCR results indicated that deletion of *dnaQ* had no impact on the induction of DNA repair genes under oxidative stress (**Fig. S4C**).

**Figure 3.**
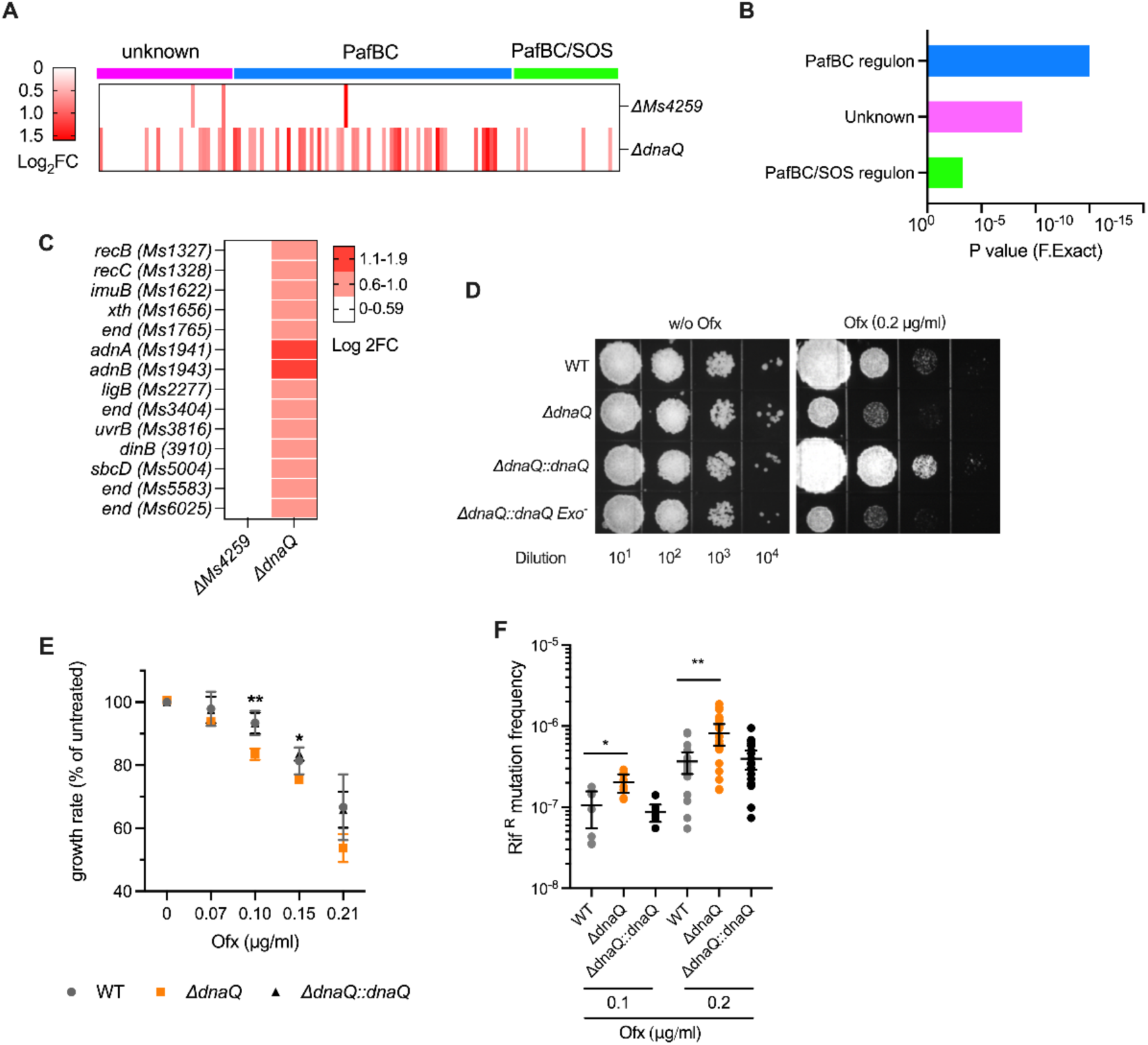
DnaQ deficiency results in replication fork perturbation and affects DNA replication fidelity upon topological stress. (A-C) transcriptional profiling of WT and the mutant strains deficient in DnaQ or DnaQ homolog encoded by *Ms4259*. (A) DnaQ-depleted cells induce a DNA damage response (DDR), DDR genes were categorized according to the established mycobacterial DDR regulons (PafBC/SOS regulon, PafBC regulon and unkonwn). (B) Enrichment analyses of the differential expressed genes in the DnaQ-depleted cells according to three DDR regulons. (C) Differential expressed PafBC regulon genes in indicated strains. End, endonuclease. (D) Growth of the indicated strains assessed by 10-fold serial dilutions on 7H10 plates with or without sub-MIC Ofx. (E) Growth rate of the indicated strains in the presence of sub-MIC of Ofx. Bacterial growth was measure by OD_600_, growth rate was calculated from exponential-phase of growth (OD_600_ within 0.1~0.4) and expressed relative to the untreated counterparts. Data shown are mean ± 95% CI. \**P*<0.05, \*\**P*<0.01 using an unpaired t test compared to WT. (F) Rif^R^ mutation frequencies of the indicated strains after exposure to Ofx. Data shown are mean ± 95% CI. \**P*<0.05, \*\**P*<0.01 using an unpaired t test on log transformed values.

Given that no growth defect was observed in the *dnaQ* mutant (**Fig.S2B**), as well as the inability of Exo^−^ DnaQ variant to restore the Rif^R^ frequency in the *ΔdnaQ* mutant (**Fig.1A**), we conclude that DnaQ depletion unlikely impairs the assembly of replisome during normal growth^9,10^. Therefore, the enriched upregulation of PafBC regulon genes in the *ΔdnaQ* mutant may likely reflect a replication conflict under certain circumstance^27,32^. To identify this condition, we examined the susceptibility of the *dnaQ^+^* prototype and the *ΔdnaQ* mutant to a panel of replication inhibitors and genotoxic agents. We observed no difference in growth or survival between WT and the *ΔdnaQ* mutant in the presence of DNA damaging agents 4-nitroquinoline-1-oxide (4NQO), UV (cause cyclobutene pyrimidine dimers, cross-links and strand breaks) and reactive oxygen species (menadione and tert-butyl hydroperoxide), the crosslinking and alkylating agents mitomycin C, or the topoisomerase IV (function in separation of two catenated circular chromosomes to terminate replication) inhibitor etoposide (**Fig. S5A-D**). However, the *ΔdnaQ* mutant exhibited attenuated growth compared with the *dnaQ^+^* prototype upon exposure to sub-minimal inhibitory concentration (MIC) of DNA gyrase A inhibitor ofloxacin (Ofx, MIC 0.3 μg/ml) (**Fig. 3D and Fig. S5E**). This growth defect can be fully restored by expression of wild-type *dnaQ,* but not of the Exo^−^ DnaQ variant, indicating that the maintenance of cell growth upon perturbations on gyrase activity relies on DnaQ’s exonuclease activity (**Fig. 3D**). Unlike the *E. coli dnaQ* mutant, our results showed that deletion of *dnaQ* in *Msm* did not affect the ofloxacin MIC as well as the survival ability upon exposure to bactericidal concentrations of ofloxacin (1.5 μg/ml)), suggesting that mycobacterial DnaQ does not implicate in the fluoroquinolone-mediated killing (**Fig. S5F-G**)^33,34^.

### Mycobacterial DnaQ sustains DNA replication fidelity upon topological stress

At low concentration of fluoroquinolone, gyrase is reversibly trapped on DNA, thereby causing the fork to stall^35,36^. Therefore, fluoroquinolone-induced topological stress may affect bacterial growth via perturbations to DNA replication, transcription or DNA damage such as SOS response^27^. Because no growth or survival defect was observed in the *ΔdnaQ* mutant upon exposure to UV and a variety of genotoxic agents (**Fig. S5A-D**), we conclude that DNA damage is unlikely to be a causative reason for the growth defect of the *ΔdnaQ* mutant upon exposed to sub-MIC Ofx. Our results also demonstrated that this growth defect is not dependent on perturbations to transcription, as no growth and survival difference was observed between the WT and the *ΔdnaQ* mutant exposed to rifampicin (**Fig. S5E**).

Previous studies found that growth of bacterial cells upon exposure to sub-MIC fluoroquinolone is largely dependent on the efficacy of DNA replication^37,38^, which could be well reflected by bacterial exponential growth rate^39^. We therefore measured the growth rate of wild-type and DnaQ-depleted strains exposed to sub-MIC of Ofx (**Fig. 3E**). Whereas both strains showed Ofx concentration-dependent inhibition of growth, the *ΔdnaQ* mutant exhibited a rate of growth ~90% that of WT in the presence of 0.33× or 0.5× MIC of Ofx (0.1 and 0.2 μg/ml) (*P*<0.05). Expression of the wild-type *dnaQ* in the mutant strain could completely restore the growth defect (**Fig. 3E**). Further, we found that, when cultivated in the presence of sub-MIC Ofx, the *ΔdnaQ* mutant exhibited increased Rif^R^ frequency over that of the WT (*P*<0.05) (**Fig. 3F**), indicating that *dnaQ* deficiency leads to increased mutagenesis upon topological stress. Considering that no growth/survival defect was observed in the *ΔdnaQ* mutant exposed to UV and chemical agents which could also lead to stalled replication forks when the replisome encounters DNA lesions^32^, these results thus support a model whereby mycobacterial DnaQ resolves fork conflict that is specifically induced by topological stress. This speculation could be further supported by the observation that *dnaQ* is the only ciprofloxacin-responsive gene that was upregulated in both *Msm* and *Mtb*^28^.

### *dnaQ* is subject to positive selection in L4.3/LAM sublineage

To assess whether the existence of an additional proofreader in mycobacteria provides an alternative avenue for *Mtb* evolution in the real-world, we analyzed the *dnaQ* sequences of globally collected 51,229 clinical *Mtb* isolates^40^. Compared with the most recent common ancestor (MRCA) DnaQ of *Mtb* complex (MTBC), we identified two lineage-defining mutations of DnaQ, with DnaQ A164V affecting L7 and L2-L4 and DnaQ D76G affecting L4 (**Fig. 4A**). While *dnaE1* was under strong purifying selection (dN/dS: 0.44), we found the selective pressure on *dnaQ* was close to neutral (dN/dS: 0.97). This could be explained as either relaxed purifying selection or a mixture of positive and negative selection on *dnaQ*. We further tested the selection on *dnaQ* in different sublineages and found that *dnaQ* was under positive selection in some sublineages while under negative selection in other sublineages (**Table S7**). The strongest positive selection was found in L4.3 sublineage that was also previously known as LAM sublineage and considered to be a generalist with worldwide distribution^41^. In total, we observed 31 mutational events of *dnaQ* in L4.3 with 26 of them being nonsynonymous (dN/dS: 1.96) (**Table S7**). Among these mutations, *dnaQ* V88A arose at early stage of L4.3 diversification and affected 16.5% (1,203/7,284) of L4.3 strains, G151R mutation also arose at mid-root position and formed a clade (**Fig. 4B and Fig. S1B**). The positive selection on *dnaQ* suggested that the mutations might have functional effects on the mutation rate of L4.3 strains. Therefore, we compared the number of SNPs accumulated in *dnaQ-*WT and *dnaQ*-mutants. Overall, *dnaQ* mutants averagely accumulated 14 more SNPs than *dnaQ*-WT strains (*P*<0.0001, Mann–Whitney U test) (**Fig. 4C**). We also found *dnaQ* V88A and G151R clades had longer tip-to-root lengths as compared to their closest neighbors without *dnaQ* mutation, further suggesting the naturally selected *dnaQ* mutants have functional effects on *Mtb* mutation rate (**Fig. 4B**).

**Figure 4.**
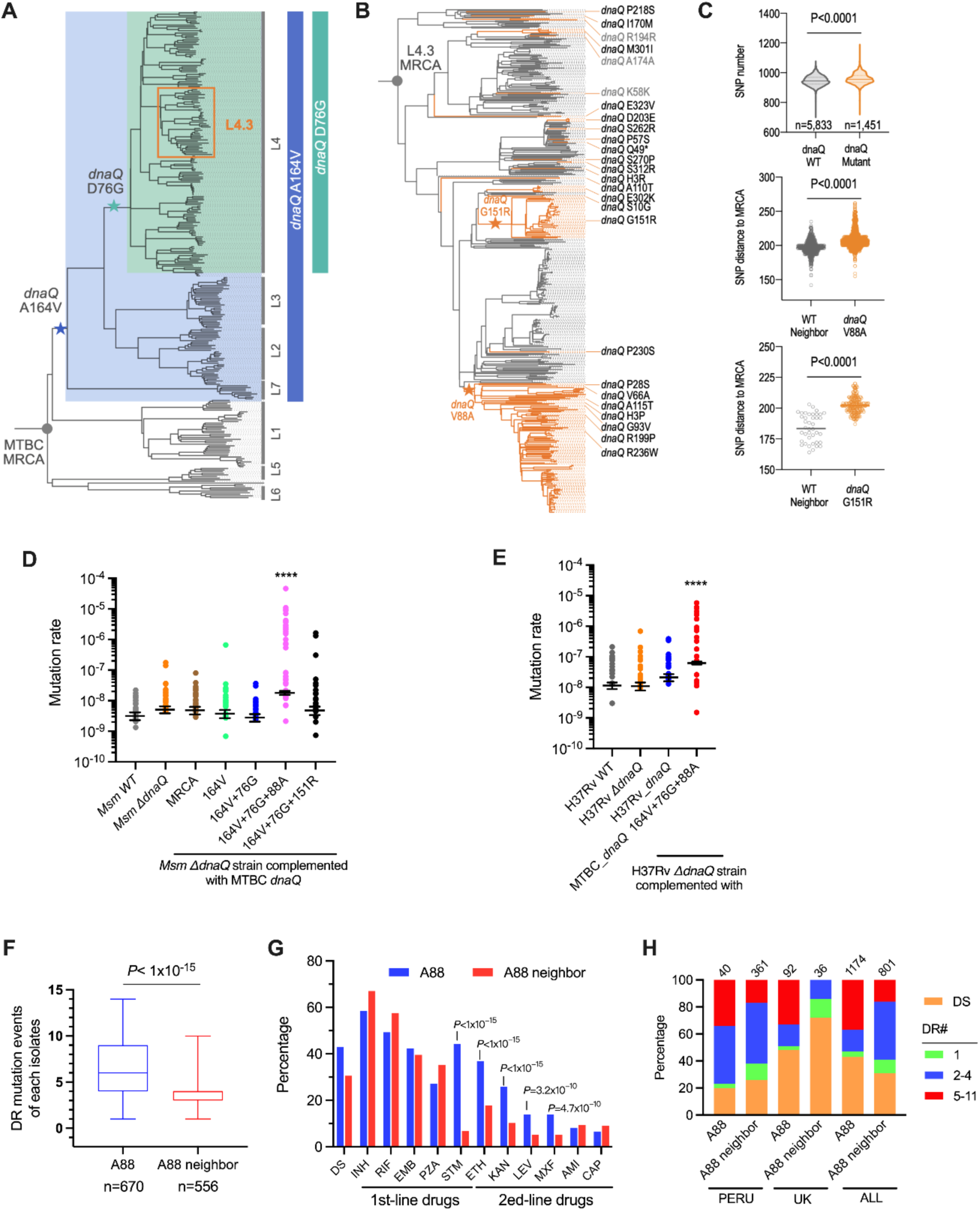
A DnaQ[V88A] variant prevalent in *Mtb* L4.3 sublineage confers hypermutability and associates with drug resistant. (A) A representative phylogenetic tree highlighting the stepwise mutations of *dnaQ* in *Mtb* lineages. (B) A phylogenetic tree of L4.3 with the mutational events of *dnaQ* marked. * refers to a pre-stop mutation. (C) Comparison of SNP numbers among *Mtb* strains within sublineage L4.3. “WT-neighbor” refer to the closest phylogenetic clades that are neighboring to *dnaQ* V88A or *dnaQ* G151R clades. (D-E) Fluctuation analyses in *Msm ΔdnaQ* mutant (D) or the *Mtb ΔdnaQ* mutant (E) expressing of a variety of *dnaQ* alleles identified in clinical *Mtb*. Circles represent mutant frequency. Estimated mutation rates were shown as mean ± 95% CI. \*\*\*\**P*<0.0001, using a one-way analysis of variance test on log transformed values compared with ancestral-type (D) or H37Rv WT (E). (F) The number of drug-resistant (DR) mutational events identified in each *Mtb* isolate containing DnaQ A88 or V88 (A88 neighbor). n indicates the number of drug resistant *Mtb* isolates. Significance was analyzed using a Mann–Whitney U test. (G) Proportion of drug resistance to individual anti-tuberculosis drug. INH, isoniazid; RIF, rifampicin; EMB, ethambutol; PZA, pyrazinamide; STM, streptomycin; ETH, ethionamide; KAN, kanamycin; LEV, levofloxacin; MXF, moxifloxacin; AMI, amikacin; CAP, capreomycin. Significance was analyzed using a Chi-square test. (H) Drug resistance profiles of *Mtb* isolates containing DnaQ A88 or V88 (A88 neighbor) in Peru, UK, and all countries (ALL). The numbers of *Mtb* isolates were indicated above the column. DS, drug sensitive. DR#, the number of drugs being resistant.

### Naturally selected *dnaQ*[V88A] enables hypermutability and is associated with extensive drug resistance

To experimentally investigate whether the naturally selected DnaQ variants would alter mutation rate, we introduced *dnaQ* mutants in the *dnaQ*-null *Msm* or *Mtb* strain H37Rv via an integrative plasmid, from which the *dnaQ* allele is expressed via a constitutive promoter. Expression of these DnaQ variants in *Msm* did not affect bacterial growth (**Fig.S2D**). By fluctuation analyses, we found the DnaQ[V88A] mutant prevalent in L4.3 caused 6-fold (*P*<0.0001) increase of the mutation rate compared with strain expressing the ancestral-type DnaQ (**Fig. 4D**), while other natural variants in the clinical isolates did not affect mutation rate. In agreement with the *Msm* results, expression of DnaQ[V88A] in the *ΔdnaQ* mutant of *Mtb* strain H37Rv also increased the mutation rate by 6-fold (*P*<0.0001) (**Fig. 4E**). These results indicated that the naturally selected *dnaQ*[V88A] is a mutator gene and leads to an intermediate potent of hypermutability^12^. Interestingly, expression of Exo^−^ DnaQ variant in the *Msm ΔdnaQ* strain only led to 2.4-fold increase of mutation rate (**Fig. 1A**), suggesting V88A’s effect might not just simply impair DnaQ exnuclease activity. Early structural and functional studies showed that the residue M85 of the *E.coli* DnaQ (corresponding to V88 of *Mtb* DnaQ) is located at *α* helix 3, a region that connects the exo I and exo II motifs and comprises the active site along with *α* helix 7 and the edges of β sheets 1–3 (**Fig. S1B**)^42,43^. However, the role of this region in the non-canonical DnaQ remains unclear.

Early studies in *E. coli*, *Salmonella* and *Pseudomonas aeruginosa* pathogens showed that the naturally selected mutators were frequently associated with inactivated MMR and multidrug resistance (MDR)^44,45^. In line with this, a mathematic modeling study in *Mtb* showed that strains with ~8-fold increased mutation rate could give rise to a notably increased risk of MDR^46^. These observations predict that the DnaQ[A88]-harboring *Mtb* isolates would be more potent to become resistant to multidrug. To test this, we compared the number of drug-resistant mutations identified in each isolate between the DnaQ[A88]-harboring population (n=1174; 670 of them contain at least one drug-resistant mutation) and the closest phylogenic neighboring isolates containing DnaQ[V88] (n=801; 556 of them contain at least one drug-resistant mutation) (**Supplementary dataset 3)**^40,47^. These isolates were distributed among 58 countries. Among the isolates containing at least one drug-resistant mutation, those containing DnaQ[A88] accumulated more drug-resistant mutations than the DnaQ[V88] isolates (mean, 6.6 *vs* 3.8; median, 6 *vs* 4) (**Fig. 4F**). Further analyses of resistance to individual drug found that the drug-resistant mutations in DnaQ[A88]-harboring isolates were associated with resistance to the second-line drugs, including levofloxacin, moxifloxacin, ethionamide and the injectable drug kanamycin (**Fig. 4G**). Accordingly, the proportion of isolates resistance to 5-11 drugs showed 2.2-fold increase in the DnaQ[A88]-harboring isolates over that of the DnaQ[V88] (36.7% *vs* 16.9%). To assess whether national medical system has an influence on these results, we further looked at *Mtb* isolates from Peru and UK, where each had >30 sequenced isolates containing DnaQ[V88] or DnaQ[A88]. Again, at both countries, the proportion of *Mtb* isolates resistant to 5-11 drugs increased at least one-fold in the DnaQ[A88] isolates than that of the DnaQ[V88] isolates (**Fig. 4H**). Together, these data provided supportive evidence that DnaQ[A88] may facilitate the *de novo* generation of drug-resistance mutations.

## Discussion

During DNA replication, a high level of fidelity is attained by the proofreading activity. In *E. coli* and other bacteria that use only one Pol III replicase (DnaE), proofreading is committed by the ε subunit 3’-5’ exonuclease DnaQ^4^. In thermophiles and mycobacteria, proofreading is mediated by an intrinsic 3’–5’ exonuclease activity posed by the PHP domain of replicative polymerase DnaE1^11,14,48^. Based on phylogenetic and structural analyses, it has speculated that proofreading in most bacteria may rely on the PHP exonuclease despite the presence of DnaQ homolog^4,11,49^. However, the biological function of this kind of alternative DnaQ exonuclease remains poorly understood. Here, we showed that the mycobacterial DnaQ is an alternative proofreader and exhibits additional role in maintenance of genomic GC content and sustaining DNA replication fidelity upon chromosome topological stress. These results demonstrate that mycobacterial replisome may deploy two proofreaders to maintain DNA replication fidelity (**Fig. 5A**).

**Figure 5.**
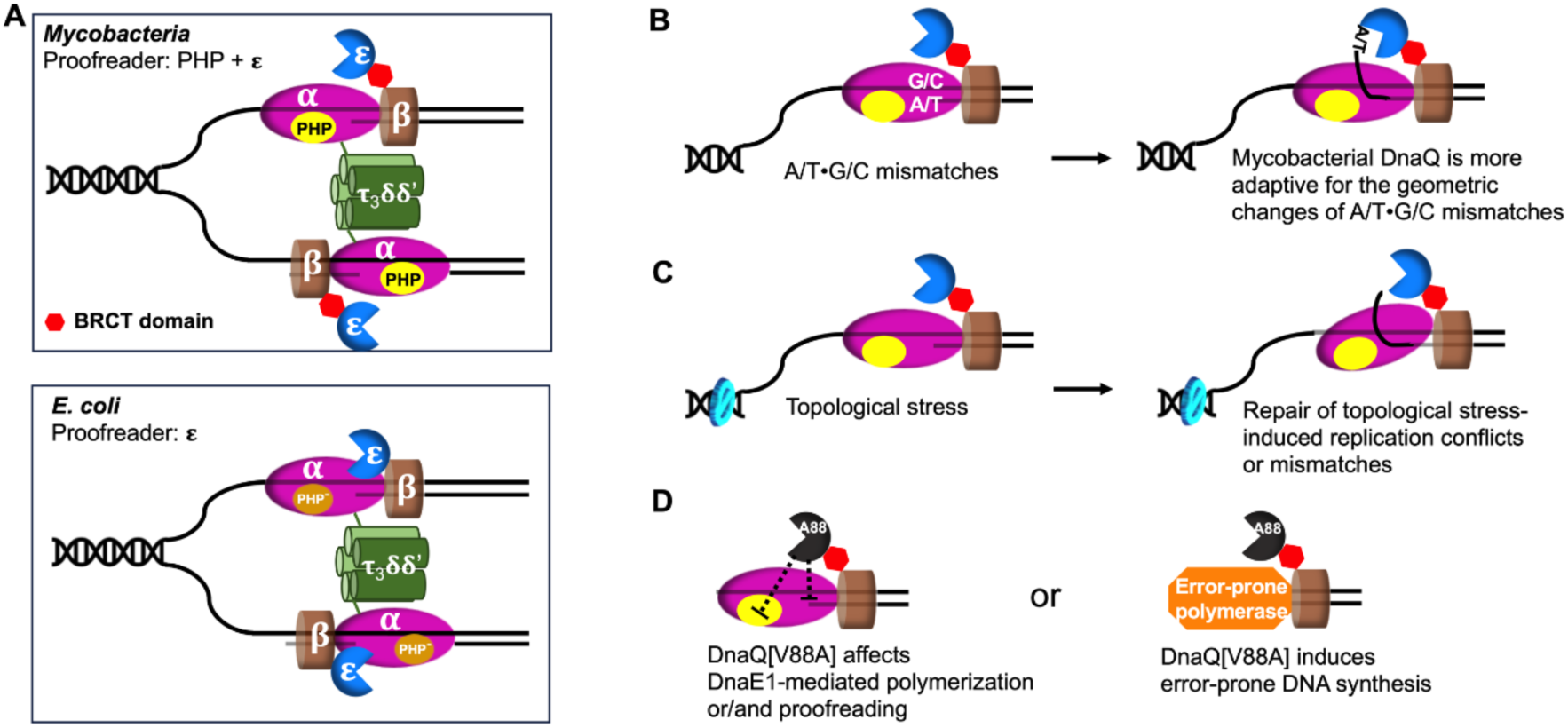
Proposed action model of DnaQ in *Mycobacteria*. (A) A schematic view of the architecture of the replisome between *Mycobacteria* and *E. coli*. The *ψ* and *χ* subunit are not shown. Compared with the *E. coli* model, the mycobacterial replisome has three distinct characters, 1) contains two proofreaders (PHP and ε subunit), 2) the ε subunit may not form a stable complex with the α subunit, and 3) contains an active PHP exonuclease domain. (B-C) Mycobacterial DnaQ participates in maintenance of genomic GC contain (B) and sustaining of DNA replication fidelity upon chromosomal topological stress (C). See also *Discussion*. (D) Possible mechanisms of DnaQ[A88]’s effect on mutagenesis.

Independent lines of evidence presented in this study indicate that mycobacterial 3’-5’ exonuclease DnaQ functions in maintaining replicative fidelity. The MA results showed that deletion of *dnaQ* led to significantly increased mutation rate, which is perceived to be largely determined by the replication fidelity of DNA polymerases^5,16,18^. Previous study deploying fluctuation analysis found that deletion of *dnaQ* in *Mtb* or in *Msm* did not significantly affect the mutation rate^11^. This discrepancy may be largely due to the difference on the methodology of detecting mutations^16–18^. The application of whole-genome sequencing to MA experiment enables complete and nearly unbiased determination of mutation events occurred in a neutral manner^16^. Further, previous study showed that expression of replicative polymerase DnaE1 with inactivated PHP exonuclease activity leads to accumulation of replicative errors in mycobacteria^11^. Therefore, the synergistical effect of DnaQ depletion and inactivation of PHP-mediated proofreading on mutagenesis signifies that DnaQ could partially rectify the replicative errors produced upon PHP inactivation. Importantly, the results of in vivo immunoprecipitation and in vitro pulldown assay collectively provide biochemical evidence that mycobacterial DnaQ binds to the replication sliding clamp via its C-terminal CBM motif, thus establishing a functional relation of DnaQ with replisome. Finally, our enzyme assays showed that the interaction with DnaN substantially stimulates DnaQ 3’-5’ exonuclease activity. Based on these data, taken together with the irrelevance of DnaQ on stress-induced mutagenesis, we conclude that mycobacterial DnaQ represents an alternative proofreading exonuclease that works together with the PHP exonuclease to maintain replicative fidelity.

Our MA data showed that, unlike the *E. coli* DnaQ, mycobacterial DnaQ exhibits additional role in maintenance of genomic GC content. How does mycobacterial DnaQ maintain GC content? Previous studies in *Salmonella* Typhimurium and *Msm* indicated that the AT-biased mutation is associated with mismatches caused by oxidized and deaminated DNA bases^50–52^. In both organisms, mutants deficient in base excision repair systems (Ung, MutY or MutM) that target oxidized and deaminated DNA bases showed strong increase of AT-biased spontaneous mutations. However, our data indicated that *dnaQ* deletion-induced AT-biased mutations may not arise from oxidative DNA damage, as exposure of the mutant to H_2_O_2_ resulted in similar mutation frequency as that of wild-type *Msm*. We propose that mycobacterial DnaQ may preferably rectify the A/T•G/C mismatches (primer•template, *i.e.* A mismatches with G or C, and T mismatches with G or C), according to the following two observations: (1) mycobacterial DnaQ specifically corrects replicative errors, and (2) deletion of *dnaQ* did not affect the A/T>G/C mutation rate. According to the structural data on thermophilic *Bacillus* high-fidelity DNA polymerase I fragment (BF) complexed with mismatched DNA, bacterial replicative DNA polymerases are more prone to extend G•T, C•T, and G•G mismatches (leading to T>C, T>G and G>C mutation), but unable to extend A/T•G/C mismatches (leading to G/C>T or G/C>A mutation) due to distortions both in the polymerase catalytic site and DNA strand (template and/or primer) geometry^53^. Because of the strong impact of A/T•G/C mismatches on geometry of template and/or primer strand, it is possible that the active sites of intramolecular exonuclease cannot efficiently adapt to this geometry change on DNA^54^ (**Fig. 5B**). This speculation could be supported by a recent cryo-EM study of the *E. coli* replisome catalytic core, showing that the T•C mismatch increases the fraying of the 3’ terminus of the primer and thus enables a ~55 Å translocation from the polymerase to the DnaQ exonuclease active site^55^. Further biochemical and structural studies are needed to fully elucidate the mechanisms underlying DnaQ’s substrate preference.

Our study also revealed a role of mycobacterial DnaQ’s 3’-5’ exonuclease activity in sustaining of DNA replication fidelity upon chromosomal topological stress. In contrast to the *E. coli* mutant deficient in *dnaQ*, which exhibited increased sensitivity to a variety of chemicals that interact with DNA or inhibit DNA synthesis^33^, our screen against a panel of genotoxic agents showed that the *ΔdnaQ* mutant *Msm* is only sensitive to sub-MIC of gyrase inhibitor Ofx. These results indicate that the *dnaQ* deletion-induced growth attenuation in the presence of sub-MIC of Ofx may stem from replication conflicts specifically induced upon topological stress (**Fig. 5C**), rather than because of impaired assembly and functioning of replisome as observed in *E. coli* strains defective in DnaQ^9,10,26^. The mild upregulation of PafBC regulon genes in the *ΔdnaQ* mutant under normal growth condition suggest that topological stress may occasionally occur in *Msm* despite the presence of a functional gyrase. In this direction, it’s noteworthy that increasing evidence shows that gyrase alone is not sufficient to resolve topological stress during bacterial replication^56,57^. For instance, it was found that GapR, a chromosome structuring protein of *Caulobacter crescentus*, participated in resolution of topological stress by regulation gyrase activity^57^.

The positive selection of *dnaQ* variations prevalent in *Mtb* L4.3/LAM sublineage provides evidence that DnaQ may play a crucial role in *Mtb’s* adaptation. Of the two clade-forming DnaQ mutations (V88A and G151R), our fluctuation analyses results demonstrated that the DnaQ[V88A] variant lead to an intermediate potent of hypermutability in both *Mtb* and *Msm*^12^. Analyses of the clinical isolates sequence data showed that the DnaQ[A88]-harboring *Mtb* accumulated more drug-resistant mutations and were more frequently associated with resistance to the second-line drugs. These results are consistent with the previous findings in the naturally evolved mutators of Gram-negative pathogens and the mathematic modeling study in *Mtb*, showing that strains with such an increased basal mutation rate could give rise to a notably increased risk of acquiring resistance to multidrug^12,44–46^. According to these results, we propose that patients infected with *Mtb* DnaQ[V88A] strain may have increased risk of treatment failures due to increased potent of this strain to acquire drug-resistant mutations. We found that the expression of DnaQ[G151R] did not results in increased mutation rate, as measured by fluctuation analysis, suggesting that the functional effect of this mutation may differ from that of V88A.

How does DnaQ[V88A] lead to increased mutagenesis? Our results showed that the expression of DnaQ[V88A] led to a notable increase of mutation rate over that of the *dnaQ*-deletion mutant or the strain expressing the DnaQ variant deficient in the 3’-5’ exonuclease activity. These results indicate that the mutational effect of DnaQ[V88A] is unlikely stemmed from depletion of DnaQ-mediated proofreading. Instead, this mutator phenotype may likely derive from perturbations on other components that ensure replicative fidelity. Considering the replisome location of DnaQ, as well as the lack of canonical MMR system in *Mycobacterium*^3^, a possible explanation is that the V88A mutation of DnaQ affects DnaE1-mediated polymerization or/and proofreading (**Fig. 5D**). This speculation could be supported by the published observations that the interaction between *E.coli* DnaQ (ε subunit) and DnaE (*α* subunit) could influence replicative fidelity: (1) suppressor mutations in *dnaE* could alleviate the growth defect and reduce the mutator phenotype of *dnaQ* mutant^58,59^, and (2) DnaQ binding enhances the stability of the clamp-*α* complex and the polymerase processivity^10,26^. Although our in vivo immunoprecipitation result did not observe an interaction between mycobacterial DnaQ and DnaE1, previous in vitro study using the analytical size exclusion chromatography showed an unstable binding between these two proteins^11^, providing evidence that mycobacterial DnaQ may interact with DnaE1 in certain circumstances. Alternatively, DnaQ[V88A]-mediated mutagenesis may rely on the action of error-prone polymerase such as DnaE2 (belong to the SOS regulon)^15,60^ (**Fig. 5D**). This model could be supported by the findings that deletion of *E. coli dnaQ* resulted in constitutive SOS response and this phenotype could be uncoupled from proofreading function^42^. Further experiments are necessary to reveal the nature of DnaQ[V88A]-mediated mutagenesis, which may promote our understanding of the dynamic regulation of mycobacterial proofreading activity.

## ACKNOWLEDGMENTS

The work was supported by the National Natural Science Foundation (grant 81991532 and 31970032 to L.-D.L., 31830002 to G.-P. Z., 31800123 to Y.-Y.X.); the Shanghai Committee of Science and Technology (grant ZD2021CY001 to L.-D.L.); the National Institutes of Health (grant P01 AI132130 to S.M.F.).

## AUTHOR CONTRIBUTIONS

Conceptualization, L.-D.L.; Methodology, L.-D.L., S.M.F., M.-Z.D. and Q.L.; Experiments, M.-Z.D., S.-J.C., F.H. Y.-Y.X. and X.C.; Clinical data analyses, Q.L. and M.Y.G.; Data analyses, M.-Z.D., Q.L., W.S. and L.-D.L.; Writing and editing, L.-D.L., Q.L., M.-Z.D. and G.-P.Z.; S.M.F. supervised the clinical data analyses; L.-D.L. supervised the whole work.

## Materials and Methods

### Strains and mutants

*M. smegmatis* mc^2^155 (ATCC 706) and *M. tuberculosis* H37Rv (ATCC 27294) were used in this study^61^. Strains and primers used in this study were listed in **Table S8** and **Table S9**, respectively. The knockout mutant strains were generated *via* allelic exchange using a specialized phage transduction method^62^. To construct double deletion mutant, the hygromycin B-resistant gene in the plasmid pYUB854 was replaced with the kanamycin-resistant gene *aph*, the resulting plasmid was used to generate the allelic exchange construct. For expression of *dnaQ* and its variants, the *dnaQ* alleles were amplified by PCR and cloned into an integrative single-copy plasmid pMV361, gene expression was controlled by a constitutive *Msm* promoter (*groL1*). For expression of DnaE1 and DanE1[D228N], DNA sequences encoding N-terminal MYC-tagged *dnaE1* and *dnaE1*[D228N] were PCR amplified and cloned into an episomal multicopy plasmid pMV261, expression was controlled by a Tet-on expression system^63^. All plasmids used in this study were verified by DNA sequencing.

### Culture conditions

Mycobacterial strains were grown in Difco Middlebrook 7H9 broth (BD #271310) or on 7H10 (BD #262710) agar with the supplementation of 0.5% glycerol, 0.05% Tween80 and 10% OADC (*Mtb*). If applicable, hygromycin B (Hyg) or kanamycin (Kan) were added to a final concentration of 50 μg/mL or 20 μg/mL, respectively. Experimental cultures were started by inoculating overnight culture into fresh media (without antibiotic) to achieve an OD_600_ of 0.01~0.02, then incubating at 37 °C with shaking at 100 rpm.

### Mutation accumulation assay

MA assay was performed as previously described^11,18^. *M. smegmatis* MA independent lines were evolved in parallel starting from the parental strain wild-type mc^2^155 and its *dnaQ*-deficient mutants (*ΔMs4259*, *ΔdnaQ*, *ΔdnaQΔMs4259*). For each genotype, 10~12 MA lines were initiate from a single colony. Each line was streaked for single colonies on a 7H10 plate (without antibiotic) and incubated for 2 days. This procedure was then followed repeatedly for the desired number of passages. The bottlenecking procedure used for this experiment ensures that mutations accumulate in an effectively neutral fashion. The number of generations (n) was then calculated by n = log_2_N, with N being the number of cells per colony. To estimate the number of cells in a generation colony, at least 10 colonies were excised from the agar plates, resuspended in PBST (PBS with 0.05% Tween 80) to generate a single-cell suspension, and dilutions were plated on 7H10 plates. For wild-type mc^2^155, *ΔMs4259* and *ΔdnaQ* strains, the average number of cells in a colony was 7.66 × 10^4^ cells, which corresponds to 16.3 generations, while the *ΔdnaQΔMs4259* strain was 3.94×10^5^ cells and corresponds to 18.6 generations.

### Whole genome sequencing analysis

Genomic DNA was isolated from 10 mL *Msm* cultures using standard CTAB extraction method. DNA concentration and purity were measured using a Qubit 3.0 fluorometer (Life Technologies) and a NanoDrop-2000 spectrophotometer (Thermo Fisher Scientific, Wilmington, DE, USA). Whole-genome sequencing libraries were constructed with the NEXTflex™ DNA Sequencing Kit compatible with the Biomek® FXp (Bio Scientific) following the manufacturer’s instructions. Sequencing was performed on the Illumina X-10 instrument to obtain 300-bp paired-end reads. The obtained Illumina reads were filtered with the Trimmomatic (version0.32) to remove low-quality bases and adapter. After that, the filtered reads were aligned to the reference genome (NC_008596) with the short-read alignment tool Bowtie2. Potential duplicates were removed with Picard tools. Mutation accumulation lines were covered to an average depth of 194× (139-360×) and 99.9% genome coverage at a depth greater than 10×. The variant calling was performed with SAMtools and Genome Analysis Toolkit^64^. A single-nucleotide polymorphism (SNP) and indels (≤30 bp) were called and filtered if (i) we have at least 10 reads covering the site, (ii) it was found at a frequency of >0.8. SNP that was observed in all lines of a strain was excluded^16^.

### Estimate of mutation rate from MA assay

The mutation rate was estimated as previously described^11,16,52^. The estimation for the mutation rate for a MA line was generated with the equation: μ = m/(N*g). m is defined by the number of variants (SNPs and indels) observed, N is determined based on >99.9% coverage of a 6,988,209 bp *Msm* mc^2^155 genome, and g is an estimate of the number of generations that occurred during passaging.

### Fluctuation analysis

Fluctuation analysis was performed as previously described^46^. For each strain, starter cultures inoculated from freezer stocks were grown to an OD_600_ of ~0.8, and then diluted by 7H9 to an OD_600_ of 0.0001 (~10,000 cells per ml). The diluted cultures were immediately split into ~30 cultures of 5 mL each in 30-mL square PETG culture bottles (Sangon Biotech) and grown at 37 °C with shaking at 100 rpm to OD_600_ of ~0.8. Cell counts were determined by plating dilutions. The cell pellet from 4.5 mL culture was plated on 7H10 agar containing 100 μg/mL rifampicin. The rifampicin-resistant colonies were counted after culturing at 37 °C for 4 d. The mutation rate was calculated by the FALCOR web tool found at https://lianglab.brocku.ca/FALCOR/ using the Ma–Sandri–Sarkar maximum likelihood method^17,65,66^.

### Mutagenesis assay

Mutagenesis assays were performed to measure the frequency of rifampicin resistance in a population. The Rif^R^ mutation frequency was calculated by dividing the number of rifampicin-resistant colonies on a rifampicin plate by the counts of the total viable cells plated. For H_2_O_2_-induced mutagenesis, cultures at OD_600_ of 0.8 were treated with 5 mM H_2_O_2_ for 2 h. For UV-induced mutagenesis, 5 mL cultures (OD~0.8) in 9-centimeter plate were exposed to 25 mJ/cm^2^ UV radiation with a Stratagene UV Stratalinker 1800. Plates were wrapped in foil to prevent potential effects of photolyase and incubated at 37 °C for 3 h. For Ofx-induced mutagenesis, cells were inoculated into 5 mL 7H9 broth containing 0.1 μg/ml or 0.2 μg/ml Ofx in 30-mL square PETG culture bottles, grown at 37 °C for 10 h. After treatment, cells were harvested by centrifugation, resuspended in 200 μL of 7H9 broth and plated on 7H10 agar plates containing 100 μg/mL rifampicin. Cell counts were determined by plating dilutions.

### RNA isolation and sequencing

*Msm* strains were grown in 10 mL 7H9 broth (in 150-mL flask) to an OD of 0.6-0.8. Cells were harvested by centrifugation (3220×g, 5 min, 25 °C). Then, cell pellet was resuspended in 1 mL TRIzol reagent (Ambion), mixed with 500 μL of 0.1mm zirconia-silicate beads. Cells were mechanically disrupted by beads beating (Bertin, Minilys) for five cycles (35 s at maximal speed) with cooling on ice for 1 min between pulses. The TRIzol-isolated RNA was treated with 10 U RNase-free DNaseI (NEB) for 30 min at 37°C, further purified using the GeneJET RNA purification kit (Thermofisher). The RNA quality was assessed using Agilent 2100 bioanalyzer. To remove rRNA, Ribo-Zero Plus rRNA Depletion Kit (Illumina) was applied according to manufacturers’ instructions. KAPA Stranded mRNA-Seq Kit Illumina® platform was used to do the library-preparation. Libraries were sequenced on Illumina HiSeq X-ten sequencing System with a read length of 150 base pairs (bps). Raw RNA-seq reads were preprocessed through trimmomatic (version 0.39)^67^. The preprocessed reads were aligned against *Msm* genes using bowtie2 (version 2.3.5.1)^68^. Gene-level read counts were summarized with samtools (version 1.9). Gene expression difference between two samples were obtained by MARS (MA-plot-based method with Random Sampling model), a package from DEGseq^69^. Genes with at least 1.5-fold change between two samples and FDR (false discovery rate) less than 0.001 were defined as differentially expressed genes.

### Quantitative real time PCR

cDNA was synthesized using the SuperScript™ IV First-Strand Synthesis System (Invitrogen) with random hexamer primer. RT-qPCR was carried out using TB Green® Premix Ex Taq™ GC (TaKaRa, RR071Q). Gene expression data were normalized to *sigA* and expressed as fold change using the 2^−ΔΔCt^ method compared to wild-type strain or untreated control.

### Sensitivity assay

For menadione sensitivity assay, cultures at exponential growth phase were diluted to an OD_600_ of 0.001 in 3.5 mL prewarmed top agar (0.6% agarose), plated on 7H10 agar. Then, a filter paper was placed on the top agar and spotted with 10 µL of 5 mM menadione. After incubation at 37 °C for 2~3 d, the diameter of the growth inhibition zone was measured. For agar-based sensitivity assays, strains were grown to exponential phase and diluted to an OD_600_ of 0.1. Serial dilutions were performed from 10^0^ to 10^−5^ in 7H9 broth, and 5 μL each dilution was spotted on 7H10 agar containing 4-nitroquinoline-1-oxide (4NQO, 1.25, 2.5, 5 μΜ), mitomycin C (8, 16, 32 nM), etoposide (25, 50, 100 μΜ), ofloxacin (0.1, 0.2, 0.4 μg/mL), or rifampicin (5, 10, 20 μg/mL). Plates were imaged after 3~4 d incubation at 37 °C. To determine MIC, strain was grown in 7H9 broth (without the supplement of antibiotic) to exponential phase, then the culture was diluted to an OD_600_ of 0.001 in 200 μL 7H9 broth containing the appropriate concentration of antibiotic. All concentrations were tested in triplicate. After cultivation at 37 °C for 3 d, cell growth was measured by OD_600_. The percentage of growth was calculated for each strain in each growth condition by normalizing the OD_600_ to that of the no-drug control.

### Killing assay

Exponential-phase cultures were diluted to an OD_600_ of 0.1 and treated with1.5 μg/mL ofloxacin at 37 °C. For treatment with tert-Butyl hydroperoxide (tBHP), cultures at OD_600_ of 0.6~0.8 were exposed to 1 mM tBHP and incubated at 37 °C. Cell counts were determined by plating dilutions. For UV exposure, serial tenfold dilutions were performed in 7H9 broth, and 5 μL of each dilution was spotted on 7H10 agar. Agar plates were exposed to indicated doses of UV radiation with a Stratagene UV Stratalinker 1800 with 254-nm UV bulbs. Plates were wrapped in foil (to prevent potential effects of photolyase) and incubated at 37 °C.

### Measure of grate rate

Experimental cultures were started by inoculating overnight culture into 20 mL fresh 7H9 broth (in a 150-mL flask) containing Ofx (0.07-0.3 μg/mL) to achieve an OD of ~0.02, then grown at 37 °C with shaking at 100 rpm. The OD_600_ value was measured every 1.5 h to obtain growth curves. The experiments were independently repeated for two times. According to previous study^70^, growth rate was defined as *μ*=2.303(lgOD2-lgOD1)/(t2-t1) and was determined at an early time (0.1<OD<0.4) when the OD value was linearly correlated with cell density. For each growth curve, we used several time intervals to calculate the growth rate.

### Immunoprecipitation

FLAG-tagged *dnaQ* under the control of native promoter was integrated at the *attB* site of the *ΔdnaQ* mutant strain. Cells were cultivated in 50 mL 7H9 broth (without antibiotic) in a 250-mL flask to an OD of ~0.8, harvested by centrifugation at 3220× g, 5 min at 4 °C. Cell pellets were washed by 10 mL ice-cold PBS and subsequently resuspended in 0.5 mL cell lysis buffer (PBS, pH=8.0, 150 mM NaCl, 0.25% NP-40, ice-cold) containing 250 μL of 0.1mm zirconia-silicate beads, lysed by bead beating (Bertin, Minilys) for five cycles (35 s at maximal speed) with cooling on ice for 1 min between pulses. The cell lysates were centrifuged at 18,000× g for 10 min at 4 °C, the supernatant was collected for immunoprecipitation. Protein concentration was determined by the Bradford method (Sigma-Aldrich, B6916). 50 μL magnetic beads (Invitrogen, 10007D) were washed and resuspended in 200 μL Ab Bind and Wash Buffer (Invitrogen, 10007D) containing 8 μg Monoclonal ANTI-FLAG® M2 antibody (Sigma Aldrich, F1804). After incubation at room temperate for 20 min, the beads were washed by 200 μL Ab Bind and Wash Buffer, then mixed with 350 μL cell lysate supernatant (2-3 mg protein) and 350 μL cell lysis buffer, incubated at 4 °C for 3 h with rolling. Then, the beads were washed 5 times each with 200 μL cell lysis buffer. Bead-bound proteins were eluted with elution buffer (100 mM glycine-HCl, pH 3.0). The elutes were analyzed by SDS-PAGE using a Fast Silver Stain Kit (Beyotime, P0017S), or store at −80 °C for LC-MS/MS analysis.

### Mass spectrometry

The eluted sample was digested by trypsin at 37 °C for 16-18 h. The peptide mixture was loaded onto a reverse phase trap column (Thermo Scientific Acclaim PepMap100, 100 μm*2cm, nanoViper C18) connected to the C18-reversed phase analytical column (Thermo Scientific Easy Column, 10 cm long, 75 μm inner diameter, 3μm resin) in buffer A (0.1% Formic acid) and separated with a linear gradient of buffer B (84% acetonitrile and 0.1% formic acid) at a flow rate of 300 nL/min controlled by IntelliFlow technology. LC-MS/MS analysis was performed on a Q Exactive mass spectrometer (Thermo Scientific) that was coupled to Easy nLC (Thermo Scientific) for 30/60/120/240 min (determined by project proposal). The mass spectrometer was operated in positive ion mode. MS data was acquired using a data-dependent top20 method dynamically choosing the most abundant precursor ions from the survey scan (300-1800 m/z) for higher energy collision dissociation (HCD) fragmentation. Automatic gain control (AGC) target was set to 1E6, maximum inject time to 50 ms, and number of scan ranges to 1. Dynamic exclusion duration was 30.0 s. Survey scans were acquired at a resolution of 70,000 at m/z 100 and resolution for HCD spectra was set to 17,500 at m/z 100, AGC target was set to 1E5, isolation width was 1.5 m/z, microscans to 1, and maximum inject time to 50 ms. Normalized collision energy was 27 eV and the underfill ratio, which specifies the minimum percentage of the target value likely to be reached at maximum fill time, was defined as 0.1%. The instrument was run with peptide recognition mode enabled. MS/MS spectra were searched using MASCOT engine (Matrix Science, London, UK; version 2.2) against a nonredundant International Protein Index arabidopsis sequence database v3.85 (released in September 2011; 39679 sequences) from the European Bioinformatics Institute (http://www.ebi.ac.uk/). For protein identification, the following options were used. Peptide mass tolerance= 20 ppm, MS/MS tolerance= 0.1 Da, Enzyme=Trypsin, Missed cleavage= 2, Fixed modification: Carbamidomethyl (C), Variable modification: Oxidation(M)

### Western Blotting

Cell lysates was prepared as described in Immunoprecipitation. Approximately 50 μg of total cell proteins were resolved by 4%–20% SurePAGE, Bis-Tris (GenScript, M00657) at 120V for 1h and detected by immunoblotting. Proteins were transferred to PVDF membrane (Millipore, IPVH00010) at 20V, 4 °C for 1 h using a Mini Gel Tank (Life Technologies, A25977), blocked with blocking buffer (20 mM Tris-HCl pH 7.6, 150 mM sodium chloride, 0.01% (w/v) Tween-20, 3% BSA) overnight at 4 °C, washed for 5 min at room temperature with TBST (20 mM Tris-HCl pH 7.6, 150 mM sodium chloride, 0.01% (w/v) Tween-20) two times. For detecting FLAG-tagged DnaQ, the membrane was incubated for 2 h at room temperature with a monoclonal anti-FLAG antibody (Sigma Aldrich, F1804) diluted 1:1,000 in blocking buffer. Mycobacterial Hsp65 was detected using a monoclonal antibody (Abnova, MAB4853) diluted 1:5,000. Membrane was washed for 5 min at room temperature with TBST five times, incubated for 1 h at room temperature with an anti-mouse IgG secondary antibody conjugated with peroxidase (Sigma Aldrich, A9044) diluted 1:10,000 in blocking buffer. Membrane was washed for 5 min at room temperature with TBST two times, blot signals were visualized using the SuperSignal™ West Pico PLUS (Thermo Scientific, 34577) and a chemiluminescence imaging system (Tanon 5200).

### Protein expression and purification

Wild-type *dnaQ*, *dnaQ*_Cdel and *dnaN* were cloned into the pET32a plasmid and transformed into the *E. coli* BL21(DE3) strain. *E. coli* cells were cultivated in LB to an OD_600_ of 0.6, expression was induced with 0.5 mM isopropyl β-D-1-thiogalactopyranoside (IPTG) at 16 °C for 12 h. Cells were harvested by centrifugation at 3220× g, 20 min at 4 °C, resuspended in buffer A (20 mM Tris-HCl, pH 8.0, 0.5 M NaCl, 10% glycerol, 10 mM imidazole and 2 mM DTT) with protease inhibitor and lysed by ultrasonication. Cell lysates were centrifuged at 18,000× g for 20 min at 4 °C to remove cell debris and insoluble fraction. The resulting supernatant was applied to a 10-mL column with 1 mL of Ni-Sepharose (GE Healthcare Life Sciences, 17531801) equilibrated with binding buffer (20 mM Tris-HCl, pH 8.0, 500 mM NaCl, 10 mM imidazole,10% glycerol, 5 mM DTT). The target proteins were eluted by 50 mM imidazole. The purified protein was concentrated to 5 mg/ml by ultrafiltration (Millipore, UFC901096) and maintained in 20 mM Tris-HCl, pH 8.0, 100 mM NaCl, 5 mM DTT, and 20% glycerol, divided into small aliquots, and stored at −80 °C. The protein concentration was determined by the Bradford method.

### Pulldown assay

To remove the N-terminal Trx_His_FLAG tag, the purified DnaN protein was digested by recombinant enterokinase (1.2 U per 50 μg DnaN) at 4 °C for 20 min, then incubated with Ni-Sepharose for 2 h at 4 °C. The elution containing native DnaN protein was concentrated by ultrafiltration (Millipore, UFC901096) and maintained in 20 mM Tris-HCl, pH 8.0, 100 mM NaCl, 5 mM DTT. To perform pulldown assay, 30 μL Ni-Sepharose was pre-bound with ~200 μg His-tagged DnaQ or DnaQ_Cdel for 5 min at 4 °C. After washing 2 times with 300 μL washing buffer (20 mM Tris-HCl, pH 8.0, 100 mM NaCl, 10% glycerol and 25 mM imidazole), the pre-bound Ni-Sepharose was mixed with ~200 μg native DnaN proteins and incubated at 4 °C for 5 min. The Ni-Sepharose was washed 2 times and eluted with SDS-PAGE sample buffer. Elutions were analyzed by SDS-PAGE using Coomassie blue staining (Beyotime, P0017).

### Exonuclease assay

The N-terminal fusion tag of the purified DnaQ, DnaQ-Cdel and DnaN proteins were remove by recombinant enterokinase as described in the pulldown assay. Protein concentration was determined by Bradford method. Exonuclease assays were performed in 20 mM Tris-HCl, pH 8.0, 100 mM NaCl, 10% glycerol, 2 mM MgCl_2_, 2 mM DTT and 50 μM BSA. Reactions were performed at 25 °C with 5 nM purified protein and 38 nM DNA substrate 5’-FAM-GTTCACGAGACCTACTGAC ACTGA-3’, and stopped by adding 100 mM EDTA, pH 7.4. The reaction products were analyzed by a denaturing 20% acrylamide gel and imaged with a Typhoon Imager (GE Healthcare).

### WGS data analysis

Whole-genome sequencing data for 51,229 *Mtb* isolates from 203 SRA projects included in this study were described previously^40^. The tool *prefetch* (https://www.ncbi.nlm.nih.gov/books/NBK242621/) was used for downloading the *sra* files of the *Mtb* isolates. The downloaded *sra* files were converted to paired-end or single-end *fastq* files using the toolkit of *fastq-dump* (https://www.ncbi.nlm.nih.gov/books/NBK158900/). Samples with an average sequencing depth over 20× and a mapping rate of over 90% were used for downstream analyses. Sequencing reads were trimmed with *Sickle*^71^. Trimmed reads with length > 30 and Phred scores > 20 were retained for subsequent analyses. The inferred ancestral genome of the most recent common ancestor of the *MTB*C was used as the reference template for reads mapping^72^. Sequencing reads were mapped to the reference genome using *Bowtie* 2 (v2.2.9)^68^. *SAMtools* (version 1.3.1) was used for SNP calling with the minimal mapping quality set to be 30^73^. We excluded SNPs located in repetitive regions of the genome (e.g., PPE/PE-PGRS family genes, pro-phage genes, insertion, or mobile genetic elements) that are difficult to characterize with short-read sequencing technologies^74^. Fixed SNPs with a frequency of ≥95% and at least 10 supporting reads were identified using *VarScan* (v2.3.9) with the strand bias filter on. Small insertions or deletions (INDELs) identified by *VarScan* (v2.3.9) were also excluded. *Mtb* isolates were typed into lineages based on the previously defined lineage-specific barcode SNPs^75,76^. We excluded isolates showing heterozygous typing results or those that were missing lineage-defining SNPs in the lineage-associated analysis.

To test whether *dnaQ* was under positive selection in *Mtb* population, we used a previously described method (pNpS) to generate mutations *in silico* through a random substitution process^77^. Briefly, a codon substitution matrix was generated using a base substitution model which considers the genome’s GC content (65.6% for *Mtb*) and the proportion of transitions that occurred at the wobble position of codons in synonymous fixed mutations (Ti, 0.729). For each codon, a simulation of 50,000 individual introductions of a single mutation was performed and the outcomes were scored as either synonymous or nonsynonymous. The average number of nonsynonymous outcomes of the simulations is an estimate of the probability that a mutation in the given codon would be nonsynonymous.

### Statistical analysis

Significance tests were performed in GraphPad Prism version 9.4.1. Normality and lognormality tests were performed for each dataset. All performed statistical tests were two sided. Mann–Whitney U test was performed for unpaired nonparametric tests, and t tests or one-way analysis of variance (ANOVA) followed by Dunnett’s multiple comparisons test were performed for unpaired parametric tests.

