## Supplemental data for "Mycobacterial DnaQ is an Alternative Proofreader Ensuring DNA Replication Fidelity"

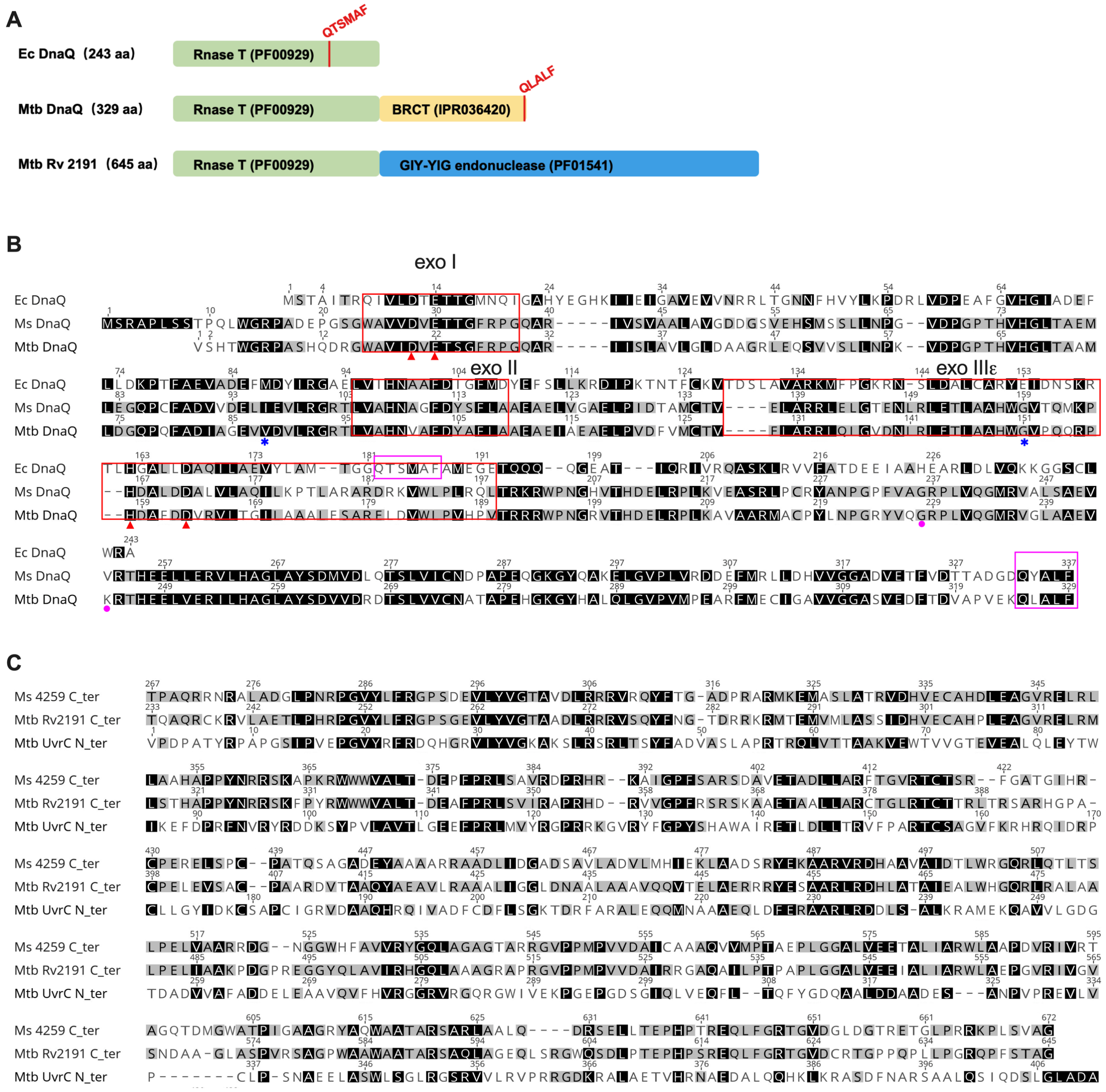


**Figure S1. Domain architectures and sequence alignments of ε exonuclease homologs.**

**A**. Domain architectures of ε exonuclease homologs from *E. coli* (*Ec*) and *M. tuberculosis* (*Mtb*). The *Mtb* DnaQ (Rv3711) was the annotated 3'–5' ε exonuclease, whereas *Mtb* Rv2191 is the second potential DnaQ homolog. Domains predicted by InterPro and the database access# were labeled, the red vertical lines and the sequences above indicate the β-clamp binding motif.

**B**. Sequence alignment of the annotated DnaQ from *E. coli* (*Ec*), *M. smegmatis* (*Msm*) and *M. tuberculosis* (*Mtb*). Three conserved exo motifs of *E. coli* DnaQ (exo I, exo II and exo IIIε) are labeled by red box[1,2]. Pink dots indicate the of *E. coli* DnaQ residues involved in the interaction with DNA polymerase α subunit. Conserved catalytic residues of *E. coli* DnaQ are indicated by red triangles below the sequences[3]. The clamp-binding motifs are boxed in pink. Residues labeled by blue asterisk presents the mutations prevalent in *Mtb* L4.3.

**C**. Sequence alignment of the endonuclease domains of the second potential DnaQ homolog (encoded by *Mtb Rv2191* and *Msm Ms4259*) and *Mtb* UvrC.

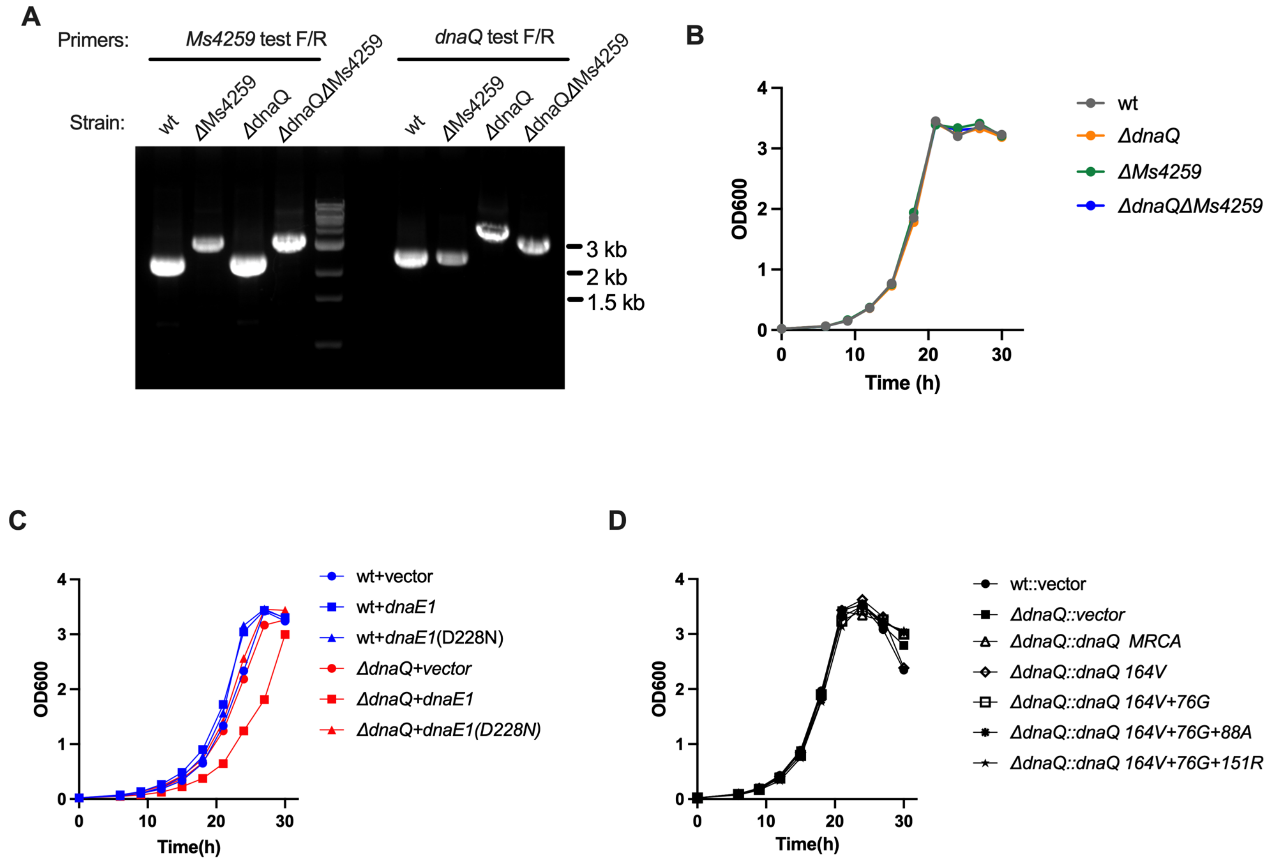


**Figure S2. Strain construction and growth curves.**

**A**. Characterization of *Msm* gene-knockout mutant strains by genotyping PCR. The primers used for genotyping PCR were designed to flank the homologous arms.

**B-D**. Growth curves of strains used for mutation rate analyses in this study. The seeding cultures were 1:200 inoculated into 7H9 and cultured at 37 °C with shaking at 100 rpm, cell growths were measured by $OD_{600}$.

A

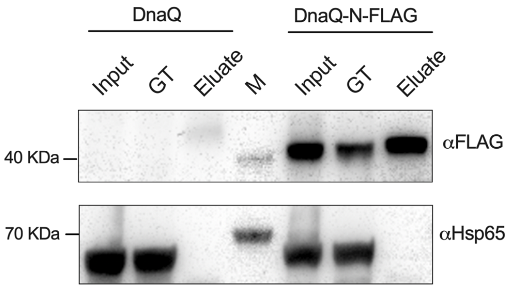


B

| Sequence | IonScore | | #PSMs |
|---|---|---|---|
| | DnaQ Co-IP | DnaQ-N-FLAG Co-IP | |
| VEISGESVVLAATDR | | 110.99 | 1 |
| QLLPAEHTAVATIGVAELTEAIKR | | 102.3 | 2 |
| VVREDFADAVAWVAR | | 85.65 | 2 |
| AGTDGNQVHLALGSGASVGK | | 72.87 | 2 |
| ALPAKPVEVSVEGTR | | 67.5 | 2 |
| QLLPAEHTAVATIGVAELTEAIK | | 59.16 | 1 |
| LSAGADDVGR | | 58.47 | 1 |
| ATTTAGLTDLK | | 57.85 | 1 |
| DGLLGIR | | 55.95 | 1 |
| VALVADR | | 55.51 | 3 |
| TDYVYLLMPVR | | 52.64 | 1 |
| TLAEAAK | | 52.59 | 1 |
| VSAEIASAGSVLVSGR | | 48.12 | 2 |
| SLPTRPTIPVLAGVLLTGTDEGLTISGFDYEVSAEVK | | 45.14 | 2 |
| VSLTCGSAR | | 43.61 | 2 |
| TDYVYLLMPVR | | 43.03 | 1 |
| LLSDITK | | 40.93 | 3 |
| LLDAEFPK | | 38.61 | 1 |
| VEISGESVVLAATDRFR | | 35.65 | 1 |

C

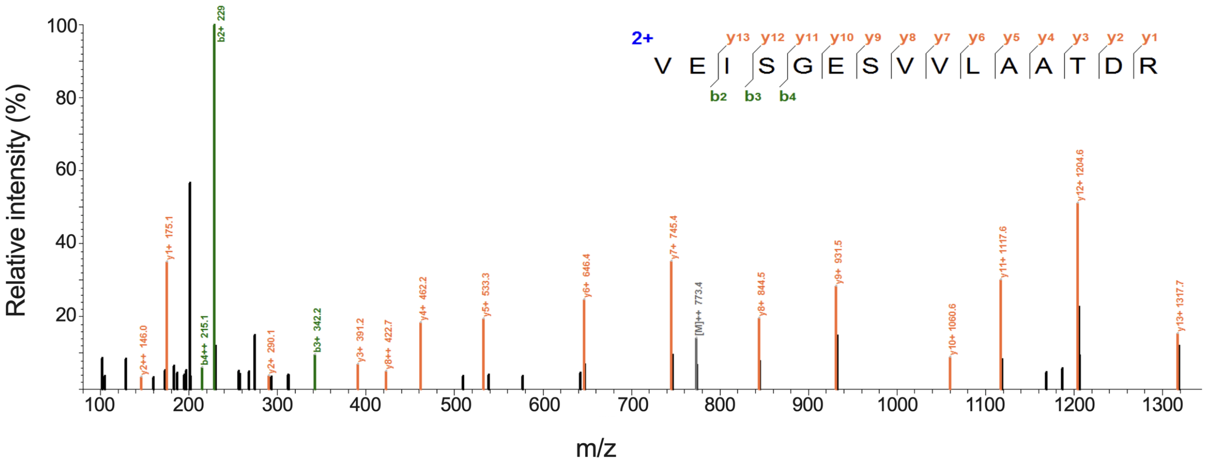


D

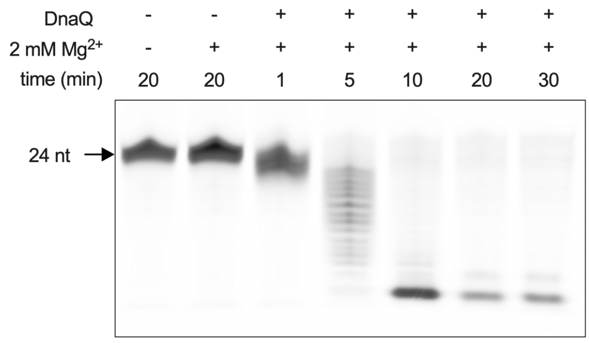


**Figure S3. DnaQ interacts directly with DnaN.**

**A**. Immunoprecipitation experiments in either *Msm ΔdnaQ*::*dnaQ* (DnaQ) or *Msm ΔdnaQ*::*dnaQ*_N-FLAG (DnaQ_N-FLAG). Inputs, go through (GT) and eluates were analyzed by western blot with FLAG antibody.

**B.** DnaN peptides identified by LC-MS/MS in the Co-IP eluates. PSMs, peptide spectrum matches.

**C**. HCD–tandem mass spectrum of a DnaN-specific peptide from Co-IP eluate in *Msm ΔdnaQ*::*dnaQ*_N-FLAG.

**D.** Time courses of DnaQ 3'-5' exonuclease activity on single-strand DNA (ssDNA). Reactions were performed with 0.19 nm purified DnaQ.

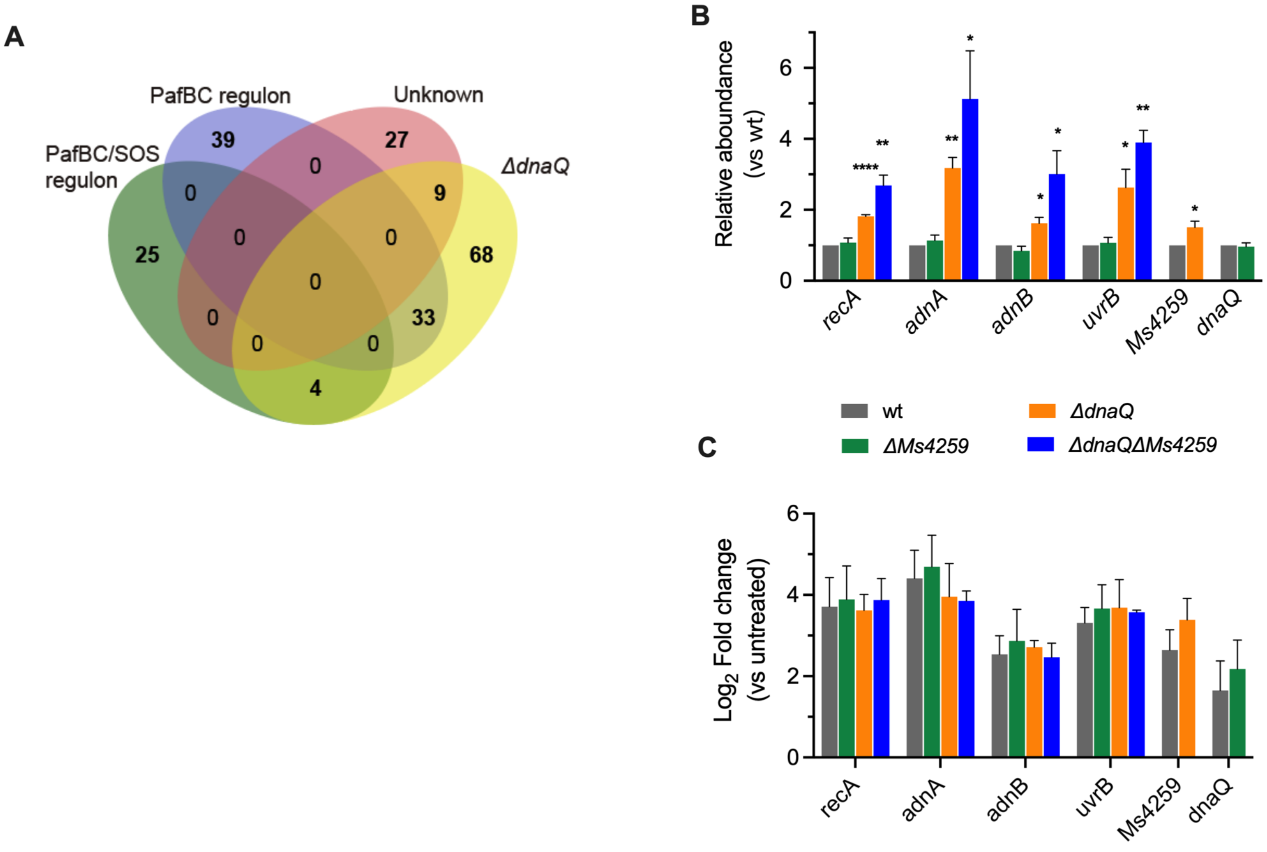


**Figure S4. Transcriptional profiling of the *ΔdnaQ* mutant strain under exponential phase of growth reveals a signature of replication fork perturbation.**

**A.** Venn diagrams categorizing of differentially expressed genes ($log_2$ fold change of ≥ 1.5, FDR< 0.001) in the *ΔdnaQ* mutant strain according to the DNA damage response regulons (PafBC/SOS regulon, PafBC regulon and unkonwn) established in *Mycobacterium* [4,5].

**B**. qRT-PCR validation of selected differentially expressed genes involved in DNA damage response. Transcript levels were normalized relative to *sigA* and expressed as fold change from wild type.

**C**. Expression of the selected genes in *Msm* exposed to $H_2O_2$ for 50 min. Transcript levels were normalized relative to *sigA* and expressed as fold change from untreated.

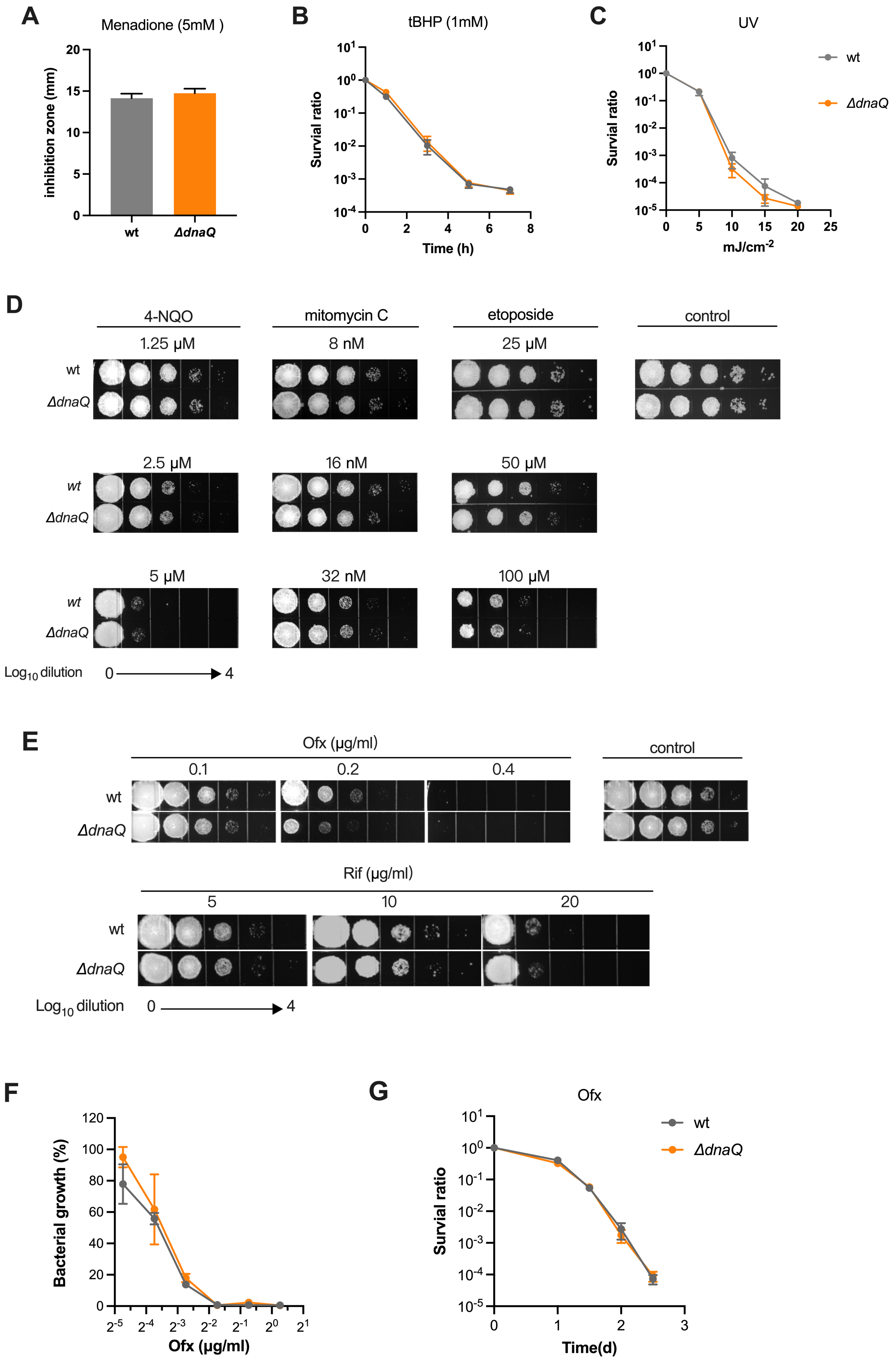
A
Menadione (5mM )
inhibition zone (mm)
wt
ΔdnaQ
B
tBHP (1mM)
Survial ratio
Time (h)
C
UV
Survial ratio
mJ/cm-2
wt
ΔdnaQ
D
4-NQO
mitomycin C
etoposide
control
1.25 μM
8 nM
25 μM
2.5 μM
16 nM
50 μM
5 μM
32 nM
100 μM
Log10 dilution 0 4
E
Ofx (μg/ml)
0.1
0.2
0.4
control
Rif (μg/ml)
5
10
20
Log10 dilution 0 4
F
Bacterial growth (%)
Ofx (μg/ml)
G
Ofx
Survial ratio
Time(d)
wt
ΔdnaQ

**Figure S5. Susceptibility of wild-type and the *ΔdnaQ* mutant strains to replication inhibitors and genotoxic agents.**

**A.** Menadine sensitivities of the indicated *Msm* strains determined by disk diffusion assay. Results show the diameter of the inhibition zone.

**B-C**. Survival ratio of the indicated *Msm* strains exposure to 1mM tert-butyl hydroperoxide (tBHP) (B) or UV (C), bacterial survival was measured by plating.

**D-E**. Sensitivities of the indicated Msm strains to genotoxic agents 4-nitroquinoline-1-oxide (4NQO), mitomycin C, etoposide (D) and ofloxacin and rifampicin (E). bacterial cultures were serially diluted and plated onto 7H10 agar plates containing indicated concentration of genotoxic agents. Plates were incubated at 37 °C for 3 days.

**F-G**. Ofloxacin susceptibility of the indicated *Msm* strains. Minimal inhibitory concentration (MIC) was determined by bacterial growth ($OD_{600}$) in the presence of ofloxacin at different concentrations (F). Antibiotic killing assay was performed by exposure of exponential cultures to 5xMIC of ofloxacin (1.5μg/ml), bacterial survival was measured by plating (G).

**Table S1. Estimated mutation rate from the fluctuation analysis data.**

| Strain | Mutation rate (per generation)$\times 10^{-9}$ | 95% confidence interval $\times 10^{-9}$ | Total cultures |
|---|---|---|---|
| WT | 3.19 | 2.31-4.17 | 30 |
| *ΔdnaQ* | 5.15 | 3.88-6.54 | 29 |
| *ΔMs4259* | 4.17 | 3.12-5.33 | 28 |
| *ΔdnaQΔMs4259* | 5.14 | 3.87-6.53 | 30 |

**Table S2. Mutational spectrum identified in MA experiment.**

| | *WT* | | *ΔMs4259* | | *ΔdnaQ* | | *ΔdnaQΔMs4259* | |
|---|---|---|---|---|---|---|---|---|
| | **Number** | **Fraction** | **Number** | **Fraction** | **Number** | **Fraction** | **Number** | **Fraction** |
| **Type of substitution** | | | | | | | | |
| **Transitions** | 13 | 0.57 | 17 | 0.53 | 34 | 0.68 | 30 | 0.68 |
| A:T>G:C | 5 | 0.22 | 8 | 0.25 | 7 | 0.14 | 6 | 0.14 |
| G:C>A:T | 8 | 0.35 | 9 | 0.28 | 27 | 0.54 | 24 | 0.55 |
| **Transversions** | 10 | 0.43 | 15 | 0.47 | 16 | 0.32 | 14 | 0.32 |
| A:T>C:G | 3 | 0.13 | 5 | 0.16 | 4 | 0.08 | 3 | 0.07 |
| G:C>C:G | 1 | 0.04 | 1 | 0.03 | 3 | 0.06 | 2 | 0.05 |
| G:C>T:A | 4 | 0.17 | 8 | 0.25 | 8 | 0.16 | 9 | 0.20 |
| A:T>T:A | 2 | 0.09 | 1 | 0.03 | 1 | 0.02 | 0 | 0 |
| A:T sites | 10 | 0.43 | 14 | 0.44 | 12 | 0.24 | 9 | 0.20 |
| G:C sites | 13 | 0.57 | 18 | 0.56 | 38 | 0.76 | 35 | 0.80 |
| Total BPSs | 23 | | 32 | | 50 | | 44 | |
| Transition/Transversion | 1.30 | | 1.13 | | 2.13 | | 1.25 | |
| **Consequences of substitutions** | | | | | | | | |
| **Position** | | | | | | | | |
| Noncoding | 6 | 0.26 | 10 | 0.31 | 3 | 0.06 | 9 | 0.2 |
| Coding | 17 | 0.74 | 22 | 0.69 | 47 | 0.94 | 35 | 0.8 |
| Coding/All | 0.74 | | 0.69 | | 0.94 | | 0.80 | |
| **Within coding sequences** | | | | | | | | |
| Synonymous | 6 | 0.35 | 3 | 0.14 | 18 | 0.38 | 11 | 0.31 |
| Nonsynonymous | 11 | 0.65 | 19 | 0.86 | 29 | 0.62 | 24 | 0.69 |
| Nonsyn/Syn | 1.83 | | 6.33 | | 1.61 | | 2.18 | |
| **Indel events** | | | | | | | | |
| Total Indels | 10 | | 7 | | 16 | | 11 | |
| Insertions | 7 | 0.70 | 4 | 0.57 | 10 | 0.63 | 6 | 0.55 |
| Deletions | 3 | 0.30 | 3 | 0.43 | 6 | 0.37 | 5 | 0.45 |
| Insertions/deletion | 2.33 | | 1.33 | | 1.67 | | 1.2 | |

**Table S3. Mutation rate of BPSs observed in MA experiment.**

| Strain | BPSs mutation rate per base pair (± 95% CI) × $10^{-10}$ | Transition mutation rate per base pair (± 95% CI) × $10^{-10}$ | G:C>A:T mutation rate per base pair (± 95% CI) × $10^{-10}$ |
|---|---|---|---|
| *WT* | 3.38 ± 1.00 | 1.91 ± 1.00 | 1.18 ± 0.87 |
| *ΔdnaQ* | 8.76 ± 4.41 | 5.97 ± 2.73 | 4.74 ± 2.58 |
| *ΔMs4259* | 5.60 ± 1.97 | 2.99 ± 1.33 | 1.58 ± 0.71 |
| *ΔdnaQΔMs4259* | 5.68 ± 2.17 | 3.85 ± 1.35 | 3.08 ± 1.10 |

**Table S4. Mutation rate normalized to the genomic GC or AT content.**

| Strain | A/T>G/C mutation rate per base pair ( ± 95% CI)×$10^{-10}$ | G/C>A/T mutation rate per base pair (95% CI)×$10^{-10}$ | Expected GC content |
|---|---|---|---|
| *WT* | 3.40 ± 2.88 | 2.69 ± 1.47 | 55.8% |
| *ΔdnaQ* | 5.61 ± 6.76 | 9.37 ±4.62 | 37.4% |
| *ΔMs4259* | 6.63 ± 4.23 | 4.55 ± 2.03 | 59.3% |
| *ΔdnaQΔMs4259* | 3.35 ± 2.14 | 6.50 ± 2.99 | 34.3% |

**Table S5. Indels mutation rate observed in MA experiment.**

| Strain | No. of Indels | Indels mutation rate (± 95% CI) | | No. of Indels at homopolymer tract (fraction) | Mutation rate of Indels at homopolymer tract (± 95% CI) | |
|---|---|---|---|---|---|---|
| | | per genome ×$10^{-4}$ | per base pair ×$10^{-10}$ | | per genome ×$10^{-4}$ | per base pair ×$10^{-11}$ |
| *WT* | 10 | 10.17 ± 8.72 | 1.48 ± 1.25 | 4 (40%) | 4.00 ± 3.75 | 6.00 ± 5.63 |
| *ΔdnaQ* | 16 | 19.70 ± 13.85 | 2.82 ± 1.98 | 10 (63%) | 12.30 ± 7.30 | 17.70 ± 10.22 |
| *ΔMs4259* | 7 | 8.60 ± 7.32 | 1.24 ± 1.03 | 3 (43%) | 3.70 ± 6.00 | 5.30 ± 8.49 |
| *ΔdnaQΔMs4259* | 11 | 10.00 ± 5.39 | 1.38 ± 0.77 | 9 (82%) | 8.17 ± 5.90 | 11.33 ± 8.40 |

**Table S6. Indel events in homopolymeric tract observed in MA experiment.**

| Strain | #Line | Genome position | Repeat sequence | Repeat time | Mutant sequence | Event |
|---|---|---|---|---|---|---|
| WT | 1 | 886312-886319 | GGGGGGGG | 8 | GGGGGGGGG | Insertion |
| | 7 | 1280798-1280806 | CCCCCCCCC | 9 | CCCCCCCCCC | Insertion |
| | 10 | 1003779- 1003783 | CCCCC | 5 | CCCCCC | Insertion |
| | 11 | 531272- 531276 | CCCCC | 5 | CCCCCC | Insertion |
| *ΔMs4259* | 5 | 3173119- 3173124 | CCCCCC | 6 | CCCCCCC | Insertion |
| | 8 | 425258- 425264 | CCCCCCC | 7 | CCCCCC | Deletion |
| | 8 | 1280798-1280806 | CCCCCCCCC | 9 | CCCCCCCCCC | Insertion |
| *ΔdnaQ* | 1 | 5795547-5795553 | CCCCCCC | 7 | CCCCCCCC | Insertion |
| | 3 | 886312-886319 | GGGGGGGG | 8 | GGGGGGGGG | Insertion |
| | 3 | 1650148-1650154 | GGGGGGG | 7 | GGGGGGGG | Insertion |
| | 5 | 3094099-3094103 | GGGGG | 5 | GGGGGG | Insertion |
| | 6 | 5765135-5765138 | TTTT | 4 | TTTTT | Insertion |
| | 7 | 1146734-1146739 | GGGGGG | 6 | GGGGGGG | Insertion |
| | 7 | 3438339-3438344 | CCCCCC | 6 | CCCCC | Deletion |
| | 9 | 1280798-1280806 | CCCCCCCCC | 9 | CCCCCCCC | Deletion |
| | 9 | 1416609-1416611 | GGG | 3 | GGGG | Insertion |
| | 10 | 5795547-5795553 | CCCCCCC | 7 | CCCCCCCC | Insertion |
| *ΔdnaQΔMs4259* | 1 | 3580020-3580027 | CCCCCCCC | 8 | CCCCCCCCC | Insertion |
| | 2 | 344011-344013 | GGG | 3 | GGGG | Insertion |
| | 3 | 1280798-1280806 | CCCCCCCCC | 9 | CCCCCCCC | Deletion |
| | 8 | 1280798-1280806 | CCCCCCCCC | 9 | CCCCCCCC | Deletion |
| | 9 | 886312-886319 | GGGGGGGG | 8 | GGGGGGGGG | Insertion |
| | 10 | 669590-669595 | CCCCCC | 6 | CCCCC | Deletion |
| | 10 | 3111341-3111345 | GGGGG | 5 | GGGG | Deletion |
| | 10 | 5959370-5959372 | CCC | 3 | CCCC | Insertion |
| | 12 | 1338491-1338496 | CCCCCC | 6 | CCCCCCC | Insertion |

**Table S7. dN/dS value of *dnaQ* in *Mycobacterium tuberculosis* sublineages.**

| Sublineages | dN/dS | Total mutation events | Nonsynonymous mutation events | Synonymous mutation events |
|---|---|---|---|---|
| L1 | 1.03 | 70 | 49 | 21 |
| L2.2 | 0.99 | 33 | 23 | 10 |
| L2.3 | 1.00 | 32 | 22 | 10 |
| L3 | 0.87 | 40 | 26 | 14 |
| L4.1 | 0.82 | 32 | 21 | 11 |
| L4.2 | 0.14 | 8 | 2 | 6 |
| L4.3 | 1.96 | 31 | 26 | 5 |
| L4.4 | 1.36 | 12 | 9 | 3 |
| L4.5 | 1.42 | 16 | 12 | 4 |
| L4.6 | 0.25 | 6 | 3 | 3 |
| L4.7 | 0.56 | 5 | 3 | 2 |
| L4.8 | 0.69 | 32 | 19 | 13 |
| L4.9 | 1.97 | 11 | 9 | 2 |

### Table S8. Bacteria strains used in this study.

| Strain | Description |
| --- | --- |
| ***M. smegmatis*** | |
| *Msm* | wild-type *Msm* strain mc$^2$155 (ATCC 706) |
| *Msm ΔdnaQ* | *Msm* wherein the *dnaQ (MSMEG_6275)* gene is replaced with *hyg* cassette |
| *Msm ΔMs4259* | *Msm* wherein the *MSMEG_4259* gene is replaced with *hyg* cassette |
| *Msm ΔdnaQΔMs4259* | *Msm ΔMs4259* wherein the *dnaQ (MSMEG_6275)* gene is replaced with *kan* cassette |
| *Msm ΔdnaQ::dnaQ$_{Msm}$* | *dnaQ*-null *Msm* with an integrative plasmid pMV361 expressing *Msm dnaQ* |
| *Msm ΔdnaQ::Ms4259* | *Ms4259*-null *Msm* with an integrative plasmid pMV361 expressing *Msm Ms4259* |
| *Msm ΔdnaQ::dnaQ$_{Msm}$N-FLAG* | *dnaQ*-null *Msm* with an integrative plasmid pMV361 expressing a N-terminal Flag-tagged *Msm dnaQ* |
| *Msm ΔdnaQ::dnaQ$_{Msm}$C-FLAG* | *dnaQ*-null *Msm* with an integrative plasmid pMV361 expressing a C-terminal Flag-tagged *Msm dnaQ* |
| *Msm ΔdnaQ::dnaQ$_{Msm}$exo-* | *dnaQ*-null *Msm* with an integrative plasmid pMV361 expressing *Msm dnaQ* with D28A/E30A/D112A mutation |
| *Msm ΔdnaQ::dnaQ$_{Mtb}$*-MRCA | *dnaQ*-null *Msm* with an integrative plasmid pMV361 expressing MRCA MTBC *dnaQ* |
| *Msm ΔdnaQ::dnaQ$_{Mtb}$*-164V | *dnaQ*-null *Msm* with an integrative plasmid pMV361 expressing MTBC *dnaQ*[164V] |
| *Msm ΔdnaQ::dnaQ$_{Mtb}$*-164V/76G | *dnaQ*-null *Msm* with an integrative plasmid pMV361 expressing MTBC *dnaQ*[164V/76G] |
| *Msm ΔdnaQ::dnaQ$_{Mtb}$*-164V/76G/88A | *dnaQ*-null *Msm* with an integrative plasmid pMV361 expressing MTBC *dnaQ*[164V/76G/88A] |
| *Msm ΔdnaQ::dnaQ$_{Mtb}$*-164V/76G/151R | *dnaQ*-null *Msm* with an integrative plasmid pMV361 expressing MTBC *dnaQ*[164V/76G/151R ] |
| *Msm::dnaE1* | *Msm* with a multi-copy plasmid pMV261 expressing *dnaE1* from *Mtb* strain H37Rv |
| *Msm::dnaE1*[D228N] | *Msm* with a multi-copy plasmid pMV261 expressing *Mtb dnaE1*D228N] |
| *Msm ΔdnaQ:: dnaE1* | *ΔdnaQ$_{Msm}$* with a multi-copy plasmid pMV261 expressing *dnaE1* from *Mtb* strain H37Rv |
| *Msm ΔdnaQ::dnaE1*[D228N] | *ΔdnaQ$_{Msm}$* with a multi-copy plasmid pMV261 expressing *Mtb dnaE1*D228N] |
| ***M. tuberculosis* strain H37Rv** | |
| H37Rv | wild-type *Mtb* strain H37Rv (ATCC 27294) |
| H37Rv *ΔdnaQ* | *Mtb* strain H37Rv wherein the *dnaQ* (Rv3711c) gene is replaced with *hyg* cassette |
| H37Rv *ΔdnaQ::dnaQ$_{Mtb}$* | *dnaQ*-null H37Rv with an integrative plasmid pMV361 expressing H37Rv *dnaQ* |
| H37Rv *ΔdnaQ::MTBC_dnaQ* [164V/76G/88A] | *dnaQ*-null H37Rv with an integrative plasmid pMV361 expressing MTBC *dnaQ*[164V/76G/88A] |
| ***E. coli BL2(DE3)*** | |
| *dnaQ*$_{Msm}$ | *BL2(DE3)* with expression plasmid pET-32a(+) expressing Msm *dnaQ* |
| *dnaQ*$_{Msm}$-Cdel | *BL2(DE3)* with expression plasmid pET-32a(+) expressing Msm *dnaQ* without C-terminal 5 amino acids |
| *dnaN*$_{Msm}$ | *BL2(DE3)* with expression plasmid pET-32a(+) expressing Msm *dnaN* |

**Table S9. Primers used in this study.**

| Primer | Sequence (5'→3') | Purpose |
|---|---|---|
| 4259up-F | TATGGTACCCAACACACTCACGACACAGTC | *Ms4259* knockout |
| 4259up-R | TCTTCTAGAGCTTCACGGTGCACAGCACG | |
| 4259down-F | ATCAAGCTTCTCACGTCACTTCCCGAACT | |
| 4259down-R | GCCACTAGTCATGGAGCGGCTGACTTTC | |
| 6275up-F | TCTGGTACCTCGCCACCGAGAACCCCGATTTG | *Msm dnaQ* knockout |
| 6275up-R | CAATCTAGAGAGGTGGTGCGCACACACGAG | |
| 6275down-F | ATAAAGCTTAAGTTCGGCGCCCACCAGTTC | |
| 6275down-R | CGGACTAGTGAGCAGAACGATCACCGAAGGC | |
| 4259testF2 | CGAGCGTGCCTCAAGTTTTC | Knockout check primer |
| 4259testR2 | CAACAACTCTGAGCGGTCCT | |
| 6275testF2 | ACTTCGAGTTGTCGGCACAT | |
| 6275testR2 | GTCTCGTCGAACCACCACTG | |
| Kan-F | CTTGACACCGGTCGTCAGTACGCGAAGAACCAC | resistance gene *aph* replacing |
| Kan-R | CGCGCTAGCTCCTTCAACTCAGCAAAAGTTCG | |
| RPro-F4 | GGGGTACCCATCTGACGGGTGTGGTGTT | Expressing promoter |
| Rpro-R | CATGCCATGGCAATTGTCTTAGCCATTGC | |
| RPro-F6 | GCTCTAGACATCTGACGGGTGTGGTGTT | |
| MS4259-F | CATGCCATGGGCTTGACCGTCCTCTCGATCG | *Ms4259* complementation |
| MS4259-R | CCCAAGCTTCTAGCCAGCGACGCTCAGT | |
| MS6275-F | CATGCCATGGGCGTGAGCCGAGCACCGCTGT | $dnaQ_{Msm}$ complementation |
| MS6275-R | CCCAAGCTTTCAGAACAGCGCGTACTGG | |
| 306F2 | TTACCGCCTTTGAGTGAGCT | Universal primer for pMV361 |
| 306R | GATGCTGACAAACGAATAGAG | |
| Rv3711c-F | CATGCCATGGTGAGCCACACCTGGGGAC | $dnaQ_{Mtb}$ complementation |
| Rv3711c-R | CCCAAGCTTTCAGAACAACGCTAACTGCTTTTCG | |
| recA-F | GAGATCGAGGGCGAGATG | *recA* qRT-PCR |
| recA-R | GGCGTAGAACTTCAGTGCCT | |
| sigA-SF | GACGACGACATCGACGAG | *sigA* qRT-PCR |
| sigA-SR | CAGCTCCACCTCTTCTTCG | |
| adnA-F | CAGGACCGAACCCGAAACC | *adnA* qRT-PCR |
| adnA-R | ATCATGTTGGGCCACAGACC | |
| adnB-F | GGTTCTCGACTGGAAGACCG | *adnB* qRT-PCR |
| adnB-R | GACCGGAACGGACATAGTGG | |
| UvrB-F | AGGACAGCTCGATCAACGAC | *UvrB* qRT-PCR |
| UvrB-R | AACGGTCCAGATACGACTGC | |
| 6275-F | CCTTGAGCGGACGCAATTC | *Ms6275* qRT-PCR |
| 6275-R | GCGCAGATCCTCAAACCCA | |
| 4259-F | GTACGAGATCCCAGGCACC | *Ms4259* qRT-PCR |
| 4259-R | CGCCGAATCGAGACGTACA | |
| TetR08-F | CGGGGTACCGAGCGCCCAATACGCAAAC | Tet-inducible TetR08 |

| | | |
|---|---|---|
| TetR08-R | TCGCAGCTGCATAACATTTCCTCCGGATCCTG | region cloning |
| Ms3178-myc-F | AACTGCAGGAGCAGAAACTCATCTCAGAAGAGGATCTGATGAGCGGCACAGACGGAC | *dnaE1*$_{Msm}$ expression |
| Ms3178-R | CGTTCGAATCAGCCGAGGCAGCCGG | |
| D228N-F | CCGCCGCTGGCCACCAACAACTGTCACTACGTCACGC | *dnaE1*$_{Msm}$*–D228N* expression |
| D228N-R | GCGTGACGTAGTGACAGTTGTTGGTGGCCAGCGGCGG | |
| 3178-overlap-F | GAAATGTTATGCAGCTGCAGGAGCAGAAACTCATCTCAGAAGAGGATCTGATGAGCGGCACAGACGGACG | |
| 3178-overlap-R | CGACATCGATAAGCTTCGAATCAGCCGAGGCAGCCGGGGC | |
| Ms.6275NF-F | CATGCCATGGACTACAAAGACGATGACGACAAGGTGAGCCGAGCACCGCTGT | *dnaQ*$_{Msm}$*–N_FLAG* expression |
| Ms.6275NF-R | CCCAAGCTTTCAGAACAGCGCGTACTGG | |
| Ms.6275CF-F | CATGCCATGGGCGTGAGCCGAGCACCGCTGT | *dnaQ*$_{Msm}$*–C_FLAG* expression |
| Ms.6275CF-R | CCCAAGCTTTCACTTGTCGTCATCGTCTTTGTAGTCGAACAGCGCGTACTGGTCAC | |
| HIS-6275F | CATGCCATGGTGAGCCGAGCACCGCTGTCGAG | *dnaQ*$_{Msm}$ *and dnaQ*$_{Msm}$*-C_del* expression in *E. coli* |
| HIS-6275R | CGCGGATCCTCAGAACAGCGCGTACTGG | |
| HIS-6275CdelR | CGCGGATCCTCAGTCACCGTCGGCGGTGGTG | |
| HIS-0001R | CGCGGATCCTCAGCCCGGAAGCCGCACCG | *dnaN*$_{Msm}$ expression in *E. coli* |
| FLAG-0001F | CATGCCATGGTGGCGACGACGACGGCTGGGC | |
| 6275A1-F | TAAGACAATTGCCATGGTGAGCCGAGCACCGCTG | *dnaQ*$_{Msm}^{exo-}$ complementation |
| 6275A1-R | CCGGAAGCCCGTGGTCGCGACAGCGACGACCGCCCAGCC | |
| 6275A2-F | CGCTGTCGCGACCACGGGCTTCCGG | |
| 6275A2-R | GGCCAGGAAGCTGTAGGCGAACCCCGCGTTGTG | |
| 6275A3-F | CGCCTACAGCTTCCTGGCCGCC | |
| 6275A3-R | TCGACATCGATAAGCTTTCAGAACAGCGCGTACT | |
| 3711A1-F | TAAGACAATTGCCATGGTGAGCCACACCTGGGGA | *dnaQ*$_{Mtb}^{exo-}$ complementation |
| 3711A1-R | CGAGGTCGCGACGGCGATGACGGCCCAACCCCG | |
| 3711A2-F | CGCCGTCGCGACCTCGGGCTTTCGG | |
| 3711A2-R | GAGAAACGCATAGGCGAACGCGACATTGTG | |
| 3711A3-F | TCGCCTATGCGTTTCTCGCTGC | |
| 3711A3-R | CTACGTCGACATCGATAAGCTTTCAGAACAACGCTAAC | |
| G76D-F | ATGCTCGATGACCAGCCACAGTTCGCCGAT | Expression of clinical DnaQ mutants |
| G76D-R | TGGCTGGTCATCGAGCATGGCGGCG | |
| V88A-F | GTGAGGTTGCCGACGTGCTGCGCGGG | |
| V88A-R | CACGTCGGCAACCTCACCGGCGAT | |
| G151R-F | CATTGGCGTGTGCCCCAGCAACG | |
| G151R-R | GGGGCACACGCCAATGCGCGGCAAGT | |
| V164A-F | GACGACGCCCGGGTATTGACCGGG | |
| V164A-R | ATACCCGGGCGTCGTCGAATGCATC | |

**Reference：**
